# Conserved Evolutionary Response to Whole Genome Duplication in Angiosperms Revealed Using High Resolution Gene Expression Profiling

**DOI:** 10.1101/2024.09.12.612700

**Authors:** J. Luis Leal, Eva Hodková, Anja Billhardt, D. Magnus Eklund, Gustaf Granath, Pilar Herrera Egoavil, Jun Chen, Pascal Milesi, Jarkko Salojärvi, Martin Lascoux

**Affiliations:** Plant Ecology and Evolution, Department of Ecology and Genetics, Uppsala University, Norbyvägen 18D, 75236 Uppsala, Sweden; Department of Ecology, Faculty of Environmental Sciences, Czech University of Life Sciences Prague, Kamýcká 129, 16500 Prague, Czech Republic; Physiology and Environmental Toxicology, Department of Organismal Biology, Uppsala University, Norbyvägen 18A, 75236 Uppsala, Sweden; College of Life Sciences, Zhejiang University, Zhejiang, 310058, Hangzhou, China; Science for Life Laboratory (SciLifeLab), Uppsala University, Uppsala, Sweden; School of Biological Sciences, Nanyang Technological University, 60 Nanyang Drive, Singapore 637551, Singapore; Organismal and Evolutionary Biology Research Program, Faculty of Biological and Environmental Sciences, and Viikki Plant Science Centre, University of Helsinki, P.O. Box 65 (Viikinkaari 1), 00014 Helsinki, Finland

**Author notes:** Correspondence should be sent to: Plant Ecology and Evolution, Department of Ecology and Genetics, Uppsala University, Norbyvägen 18D, 75236 Uppsala, Sweden.

## Abstract

Autopolyploidy, the result of genome duplication within a single species, is widespread among plant lineages and believed to have played a major role in angiosperm evolution and diversification. Whole genome duplication often triggers significant morphological and ecological changes in autopolyploids vis-a-vis their diploid progenitors, which are induced by subtle changes in gene expression patterns, often of a stochastic nature. Recent results have nonetheless identified specific changes in meiotic, metabolic, and defense response pathways that seem to be commonly shared among autopolyploid species, hinting at convergent evolution. Notably, a set of 12 core meiotic genes, including several genes involved in meiotic crossover formation, has been found to undergo strong selective pressure in the aftermath of autopolyploidization. For the most part these findings have been based on the study of *Arabidopsis arenosa* and *A. lyrata* autotetraploids and the question has remained as to whether the evolutionary forces shaping the establishment and evolution of autopolyploidy in the Arabidopsis model system extend more broadly across angiosperms, an area where our knowledge is still limited. In order to address these questions, we conducted a comparative transcriptome analysis of *Betula pubescens*, a highly introgressed autotetraploid, and its diploid sister species, *B. pendula*, two birch species belonging to the Fagales order that diverged from Brassicales 120-140 Mya. Our results reveal significant changes in the expression patterns of *B. pubescens* in genes involved in secondary metabolic processes and the regulation of stress response to pathogens, in agreement with results obtained in other autopolyploid plant complexes. Allele-specific expression analysis identified 16 meiotic genes in *B. pubescens* with constrained expression patterns, strongly favoring alleles introgressed from *B. humilis* or *B. nana*, a set that includes 8 meiotic genes − *ASY1*, *ASY3*, *PDS5B*, *PRD3*, *SYN1*, *SMC3, SHOC1* and *SCC4* − previously found to be under selection in Arabidopsis autopolyploids. These results provide support to the hypothesis that whole genome duplication triggers similar genomic responses across flowering plants, and that the evolutionary path available to autopolyploids for regaining meiotic stability is highly conserved and dependent on a small group of core meiotic genes.

## Introduction

Polyploidization, the process giving rise to organisms containing more than two complete sets of chromosomes, has been observed across all eukaryotic lineages, most prominently in plants (Van de Peer et al. 2017), and has been implicated in angiosperm and vertebrate evolution and diversification (Dehal and Boore 2005, Soltis et al. 2009), namely through sub-and neo-functionalization of duplicated genes (Adams and Wendel 2005, Otto 2007, Van de Peer et al. 2017). Whole genome duplication (WGD) within a single species (autopolyploidization) is often associated with significant morphological, metabolic, and ecological changes in autopolyploids vis-a-vis their (originally isogenic) diploid progenitors (Soltis et al. 2004, Otto 2007, Parisod et al. 2010, te Beest et al. 2012, Lavania et al. 2012, Ramsey and Ramsey 2014, Soltis et al. 2016, Van de Peer et al. 2017), in part due to changes in gene expression patterns and epigenetic remodeling (Comai 2005, Song and Chen 2015). Early studies on the transcriptomic consequences of autopolyploidization in plant systems, based on spontaneous 4X *Arabidopsis thaliana* lines (Wang et al. 2006, Yu et al. 2010) or newly synthesized autopolyploid lines in *A. thaliana* (Pignatta et al. 2010), potato (*Solanum*, Stupar et al. 2007), maize (*Zea mays*, Riddle et al. 2010), cabbage (*Brassica oleracea*, Albertin et al. 2005), and the grass *Paspalum notatum* (Martelotto et al. 2005), showed that transcriptomic changes in the immediate aftermath of autopolyploidization tend to be relatively subdued and can be highly variable among strains and ecotypes (see also Doyle et al. 2008, Parisod et al. 2010, Bomblies and Madlung 2014, Fasano et al. 2016). This contrasts with results obtained in allopolyploids (the result of hybridization between two divergent species in combination with genome duplication) where pronounced changes in gene expression patterns in the aftermath of polyploidization have been documented (Wang et al. 2006, Otto 2007, del Pozo and Ramirez-Parra 2015, Spoelhof et al. 2017, Doyle and Coate 2019, Duan et al. 2023).

A more detailed analysis of differentially expressed genes, however, has revealed that although transcriptomic changes observed in autopolyploids have a stochastic nature and are often subtle, they tend to affect a very specific set of biological functions, hinting at the presence of conserved elements in the transcriptional response induced by autopolyploidization (Doyle and Coate 2019). Evidence collected from autotetraploid lines in *A. thaliana* (Yu et al. 2010, Del Pozo and Ramirez-Parra 2014), *Brassica rapa* (Braynen et al. 2017, Wang et al. 2018), *Isatis indigotica* (Lu et al. 2006, Zhou et al. 2015), *Betula platyphylla* (Mu et al. 2012), *Morus alba* (Dai et al. 2015), and established natural autopolyploids such as sea barley (*Hordeum marinum*, Zhou et al. 2019), point to significant changes in gene expression symptomatic with a major reconfiguration of metabolic pathways, transcription/epigenetic regulation, defense response against pathogens, phytohormone signaling, and reproductive processes.

Additional evidence that autopolyploidization disproportionately affects specific functional categories was obtained from genome scans for signatures of selective sweeps carried out in natural *A. arenosa*, *A. lyrata,* and *Cardamine amara* autotetraploid-diploid complexes (Yant et al. 2013, Marburger et al. 2019, Bohutínská et al. 2021a). Besides being enriched for several of the functional groups identified above (eg. metabolic processes), in all three autopolyploid systems the set of loci under selection also contained several genes implicated in genome maintenance and stability, including chromosome organization, meiosis, and DNA repair (Yant et al. 2013, Marburger et al. 2019, Bohutínská et al. 2021a). There is ample evidence that overcoming the disruptive effect of genome duplication on meiosis, in order to regain meiotic stability, is one of the key challenges faced by novel autopolyploids (Comai 2005, Cifuentes et al. 2010, Bomblies and Madlung 2014, Bomblies et al. 2015). This is illustrated by the increase in the number of genes involved in the regulation of meiotic crossover formation and dissolution found to be under selective pressure in polyploid lineages (vis-a-vis their diploid progenitors, Yant et al. 2013, Wright et al. 2015, Marburger et al. 2019, Monnahan et al. 2019, Bohutínská et al. 2021b), and by observations that chiasma frequency, and therefore the rate of meiotic crossovers, is generally lower in established autopolyploids than in their diploid counterparts (Yant et al. 2013, Bomblies 2023). Interestingly, recent results show that improved meiotic stability in 4X *A. lyrata*, a relatively young autotetraploid lineage, is observed in specimens containing several adaptive alleles introgressed from 4X *A. arenosa* (Marburger et al. 2019, Seear et al. 2020), a much older autopolyploid species, suggesting that for some plant systems the successful establishment of a natural autopolyploid population may hinge on their ability to borrow specific meiotic alleles from a (non-parental) third-species species via introgression.

These observations have raised the question of whether the set of meiotic genes under selective pressure might be universally conserved across autopolyploid plant species, implying that the range of evolutionary trajectories open to polyploid plant systems as they wrestle to regain genomic integrity might be rather limited, and that some of the cellular processes required to attain meiotic stability in the aftermath of WGD could be under convergent evolution (Marburger et al. 2019, Bohutínská et al. 2021a). This hypothesis, however, has been tested only within the Arabidopsis genus and in *Cardamine amara* (Marburger et al. 2019, Bohutínská et al. 2021a), a Brassicaceae species 17 My diverged from Arabidopsis (Huang et al. 2020). Results obtained so far paint a nuanced picture. While there is good agreement between the set of meiotic genes under selection detected in the autotetraploids *A. arenosa* and *A. lyrata* (all eight loci detected in *A. lyrata* are among those (12) found in *A. arenosa*, Yant et al. 2013, Wright et al. 2015, Marburger et al. 2019, Monnahan et al. 2019, Bohutínská et al. 2021b), the set of genes under selection detected in *C. amara* generally differs from those previously observed in Arabidopsis, although selection signals were inconclusive for several meiotic genes (Bohutínská et al. 2021a).

Here, we perform the transcriptomic analysis of *Betula pubescen*s (downy birch), an autotetraploid (Leal et al. 2024) tree species widespread in northern Eurasia and whose genome has been profoundly reshaped by extensive introgressive hybridization with *B. pendula*, *B. nana*, and *B. humilis* (Anamthawat-Jónsson and Tomasson 1990, Thórsson et al. 2001, Palmé et al. 2004, Jadwiszczak et al. 2012, Wang et al. 2014, Eidesen et al. 2015, Zohren et al. 2016, Tsuda et al. 2017, Leal et al. 2024). Since Arabidopsis (Brassicales) and Betula (Fagales) are estimated to have diverged 120-140 Mya (Huang et al. 2020, Zuntini et al. 2024), comparative gene expression profiling of *B. pubescens* and its closest extant diploid relative in western Eurasia, *B. pendula* (silver birch) will allow us to assess whether the evolutionary forces shaping the establishment and evolution of autopolyploids in the Arabidopsis model system extend more broadly across other polyploid angiosperms, an area where our knowledge is still rather limited (Soltis et al. 2016, Bomblies 2023). Our results reveal significant differences in expression patterns between 4X *B. pubescens* and 2X *B. pendula* in genes involved in secondary metabolic processes and biotic stress response, in agreement with results obtained in other autopolyploid plant complexes. Allele-specific expression analysis identified 16 meiotic genes in *B. pubescens* whose expression is constrained, for instance, strongly favoring alleles of *B. humilis* and/or *B. nana* origin, a set that includes 8 meiotic genes (*ASY1*, *ASY3*, *PDS5B*, *PRD3, SYN1, SMC3*, *SHOC1* and *SCC4*) previously found to be under strong selective pressure in Arabidopsis autopolyploids. Taken together, these results suggest that the molecular mechanisms underpinning meiotic stability in Arabidopsis autotetraploids extend more broadly across flowering plants, providing support to the hypothesis raised by Bohutínská and colleagues (2021a) that WGD triggers directional selection, and thus evolutionary convergence, on a very specific set of meiotic orthologous genes.

## Results and discussion

### Abiotic Stress Triggers Similar Physiological Responses in Diploids and Tetraploids

Seedlings of autotetraploid *B. pubescens* and its diploid sister species, *B. pendula*, were grown in a controlled environment from seeds collected from natural populations in Scandinavia (Fig. 1a) and subsequently subjected to four different temperature and water stress scenarios (Fig. 1b). For each abiotic condition, physiological, hormone, and gene expression data was collected from nine specimens per species, representing three ecotypes (three replicates per ecotype; Fig. 1b). *B. pendula* ecotypes are associated to the two main climate zones in Scandinavia, including their contact zone (Supplementary Table S1). Besides covering different climate zones, *B. pubescens* ecotypes also display significant variation in levels of introgression from *B. nana* and *B. humilis* (inset barplot in Figure 1a), with the genome in individuals from the northernmost population having been estimated to contain alleles of *B. nana* origin in ∼70% of all loci (∼30% of loci contain alleles of *B. humilis* origin; Leal et al. 2024).

**Figure 1.**
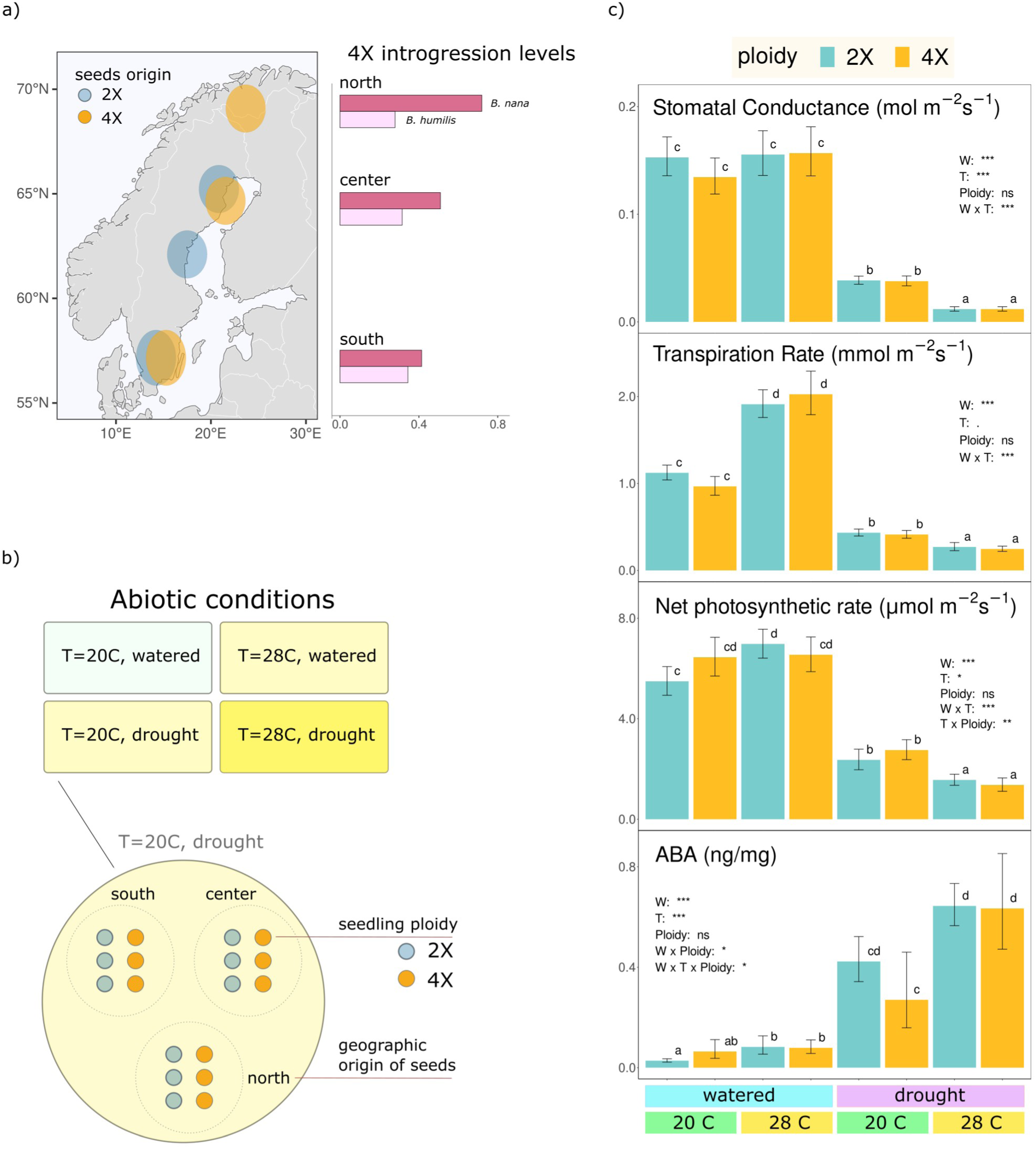
| Experimental conditions and physiological response to abiotic stress – **(a)** *B. pubescens* (4X) and *B. pendula* (2X) sampling locations in Scandinavia. For each ploidy, location of the ‘north’ ecotype coincides approximately with the species’ northernmost range limit. Barplot shows estimated fraction of the *B. pubescens* genome containing at least one allele of *B. nana* or *B. humilis* origin, obtained via introgression, based on Leal et al. (2024). (**b**) Abiotic stress conditions. For each scenario, RNA was collected from leaf and root tissue from nine diploid and tetraploid seedlings, representing all three ecotypes. (**c**) Stomatal conductance, transpiration rate, net photosynthetic rate, and levels of abscisic acid (ABA) measure in leaves as a function of ploidy, temperature, and watering conditions. Bar high indicates geometric mean for nine replicates; error bars show 95% confidence interval of the mean. Symbols above bars show results of multi-way ANOVA significance test based on the best linear model, corrected for multiple pairwise comparisons using the Tukey’s HSD test. Parameter significance values (best linear model): ‘***’: p-value < 0.001; ‘**’: p-value < 0.01; ‘*’: p-value < 0.05; ‘ns’: p-value > 0.05.

The physiological response to abiotic stress in diploid and tetraploid specimens was investigated using quantitative measures of stomatal function and photosynthesis. Stomatal conductance, a measure of stomatal aperture, decreased under drought conditions, leading to a decrease in transpiration rates and net photosynthetic rate (Fig. 1c). The combined effect of drought and high temperature produced the most severe stress scenario, inducing stomata to close in order to prevent excessive water loss via transpiration, further limiting photosynthesis. Linear model (LM) analysis confirmed that changes in all three measures were driven primarily by watering and temperature conditions (significance levels displayed in Figure 1c). Differences between ploidy levels were not statistically significant, although the LM analysis of net photosynthesis data suggests that ploidy might act as an interaction factor. Likewise, there were no obvious differences across geographic ecotypes (Supplementary Fig. S1a-c). Mean values for stomatal conductance and net photosynthetic rate observed in well-watered plants are in-line with values observed in *B. platyphylla* leaves of similar age (Hoshika et al. 2013). Leaf temperature was below ambient temperature in well-watered plants both at 20C and 28C (Supplementary Fig. S2), with transpiration rates increasing at high temperatures (Fig. 1c) to sustain efficient evaporative cooling, but overall there were no significant differences between diploid and tetraploid populations. Stomatal response to abiotic stress is mediated by the plant hormone abscisic acid (ABA; Osakabe et al. 2014, Kuromori et al. 2018). As expected, ABA measured in leaves increased dramatically under water-stress and reached maximum values when plants were subjected to both drought and high temperatures, but differences in ABA levels across ploidies were not statistically significant (Fig. 1c).

The results shown above suggest that *B. pubescens* and *B. pendula* specimens subjected to similar abiotic conditions have similar physiological responses, despite differences in ploidy, at least for the set of traits and abiotic conditions here discussed. This is congruent with findings from a comparative study of diploid and tetraploid *A. arenosa* populations which revealed no ploidy effect on a large set of morphological traits (Wos et al. 2019). Similarly, no significant differences in net photosynthetic rate were observed between diploid and autopolyploid *Medicago sativa* (likewise for *Pennisetum americanum*, Warner and Edwards 1993). In *Citrus limoni* (Rangpur lime), no differences were found in ABA levels between diploid and autotetraploid lines under water deficit, despite tetraploids having higher drought resistance (Allario et al. 2013). WGD is strongly correlated with an increase in cell size and nuclear volume, and it is known to impact important plant traits (Otto 2007, te Beest et al. 2012, Lavania et al. 2012, Soltis et al. 2016), such as inducing larger flowers, seeds, and stomata in autopolyploid *A. thaliana* lines (Del Pozo and Ramirez-Parra 2014, 2015). However, there is evidence that the increase in cell size in polyploids is often partially compensated by limiting cell division as a way to regulate organ size (Robinson et al. 2018), and it has been speculated that, depending on the trait, such compensatory mechanism could be responsible for the lack of differentiation between cytotypes (Warner and Edwards 1993, Del Pozo and Ramirez-Parra 2015, Wos et al. 2019). Also relevant in this context is the substantial level of gene flow from *B. pendula* into *B. pubescens* (Palmé et al. 2004, Wang et al. 2014, Zohren et al. 2016, Tsuda et al. 2017, Leal et al. 2024). It has been estimated that up to 80% of the genome of *B. pubescens* specimens collected in Scandinavia contain at least one allele of *B. pendula* origin (Leal et al. 2024), and such extensive levels of introgression could provide an additional rational for the presence of shared traits between two species that are often difficult to tell apart based solely on morphology (Ashburner and McAllister 2013, p. 232).

### Global Gene Expression Patterns are Primarily Determined by Tissue Type and Water Stress Conditions in Both Diploids and Tetraploids

Transcriptome analysis of leaf and root samples collected from diploid (*B. pendula*) and tetraploid (*B. pubescens*) individuals subjected to different abiotic scenarios revealed that gene expression patterns are dominated by differences across plant tissues, followed by watering conditions (Fig. 2a and Supplementary Fig. S3). Differential gene expression analysis for different contrasts conformed to these findings (Supplementary Fig. S4), with the number of differentially expressed genes (DEG) being highest for comparisons across plant tissues (8,150 (28.2%) to 10,658 (36.9%) out of 28,853 annotated genes), followed by contrasts across watering conditions in root tissue (4,953 (17.2%) to 6,666 (23.1%) genes). Analysis across ploidy levels identified between 679 (2.4 %) and 1,827 (6.3%) genes whose expression varies significantly, depending on tissue type and abiotic conditions (Fig. 2b). These values are in line with results obtained in *A. thaliana* based on cDNA microarray analysis of diploid and autotetraploid specimens, which identified between 9 and 1,589 genes whose expression varied significantly across ploidies, with the number of DEG varying widely across different natural accessions (Wang et al. 2006, Yu et al. 2010), while being generally higher in newly synthesized autotetraploid lines (Pignatta et al. 2010, Del Pozo and Ramirez-Parra 2014). Subsequent studies carried out in other diploid-autopolyploid plant systems, for the most part based on synthetic autotetraploid lines, have provided further evidence that autopolyploidization induces changes in expression affecting only a modest fraction of the genome, seldom exceeding 10% of all coding genes (Martelotto et al. 2005, Lu et al. 2006, Stupar et al. 2007, Riddle et al. 2010, Mu et al. 2012, Dai et al. 2015, Zhou et al. 2015, Fasano et al. 2016, Braynen et al. 2017, Zhou et al. 2019), although care must be exercised when comparing different studies as methodologies used to identify genes differentially expressed often relied on different experimental and statistical frameworks as well as different p-value and log2-fold change cut-off thresholds.

**Figure 2.**
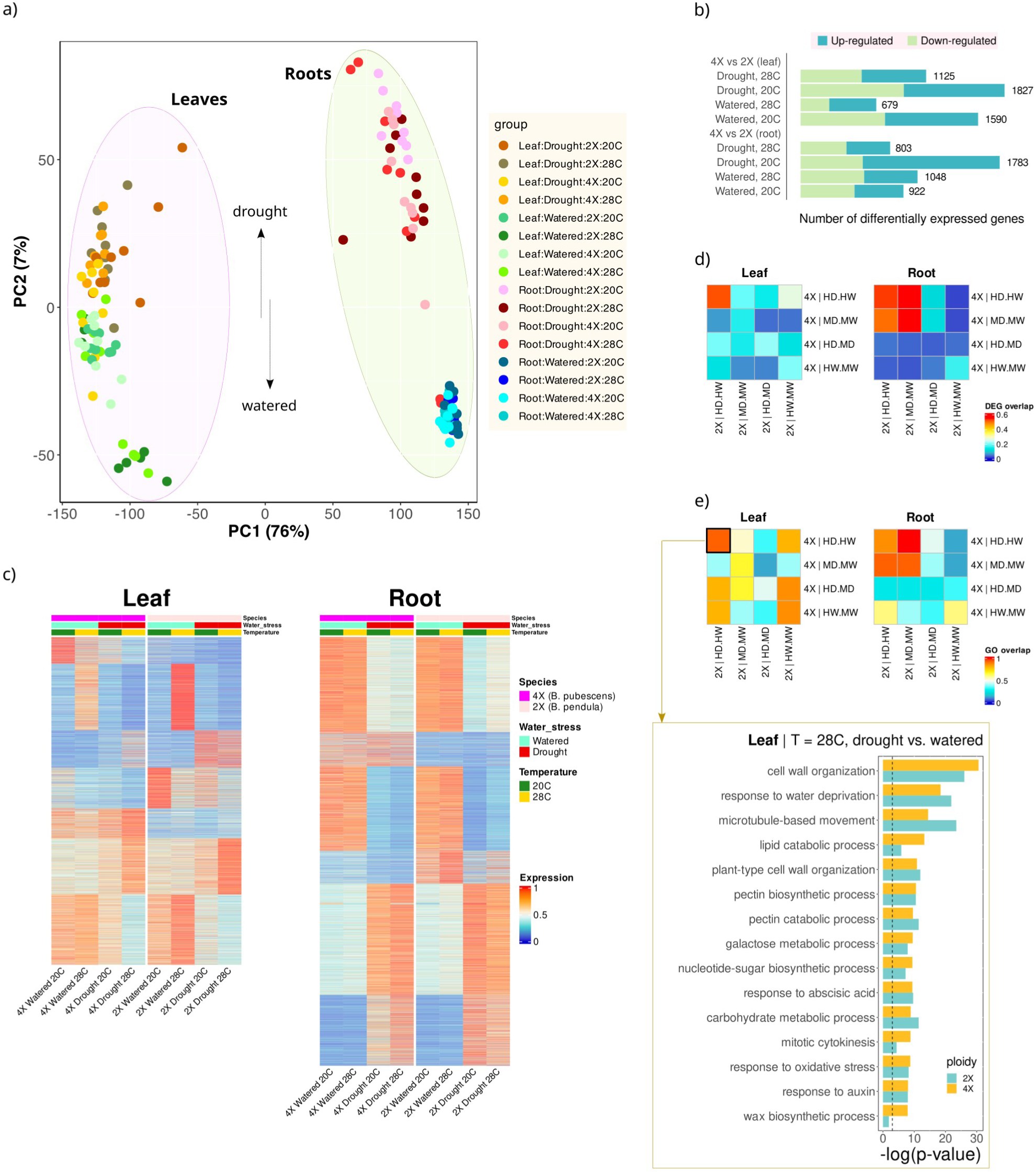
| Transcriptomic response to abiotic stress in diploids and tetraploids – (**a**) Principal component analysis of the normalized RNA-Seq count matrix for 22,519 expressed genes. Two root samples, taken from specimens kept under drought conditions, group with well-watered samples and were excluded from downstream analysis. (**b**) Number of differential expressed genes (DEG) across ploidies (*B. pubescens* (4X); *B. pendula* (2X)). Only genes with p-adj < 0.05, |log2FoldChange| > 1, and baseMean > 10 were considered to be differentially expressed. (**c**) Heatmap of normalized expression levels for 7,886 (leaf) and 10,314 (root) differentially expressed genes (rows) across different abiotic conditions and ploidies (columns). Values for each cell are the average expression for nine biological replicates, apart from *B. pubescens* root samples subjected to drought at T=28C (seven replicates). (**d**) Heatmaps showing fraction of DEG detected in *tetraploids*, across abiotic treatments, that are also listed as significant in *diploid* contrasts. ‘D’, ‘W’, ‘H’, and ‘M’ stand for ‘drought’, ‘watered’, ‘hot(28C)’, and ‘mild(20C)’, respectively. (**e**) Heatmaps showing fraction of the top-15 most significant GO categories in tetraploids, across abiotic treatments, that are also listed as significant GO terms in diploid contrasts. Barplot in inset shows the top most enriched GO categories observed in tetraploids for the set of differentially expressed genes across water treatments at 28C. Only GO categories containing at least 10 annotated genes were considered. Vertical dashed line indicates -log(p-value=0.05) threshold.

Differences between birch cytotypes became more apparent when the analysis was performed separately for each plant tissue (Fig. 2c): although 2X and 4X expression patterns have a striking resemblance, a closer look revealed some significant differences between diploids and tetraploids, both in leaves and in roots. Interestingly, cross-ploidy transcriptomic differences varied considerably depending on abiotic conditions. When we checked whether DEG identified in tetraploids, for a specific abiotic contrast, were also listed as significant for the same contrast in diploids, there was a strong overlap between gene lists (∼0.6) for contrasts across water treatments, in particular in roots (Figs. 2d), while being significantly lower for other contrasts (< 0.2), signaling a higher degree of transcriptomic divergence between cytotypes. Gene ontology (GO) enrichment analysis of differentially expressed genes revealed that the top functional categories enriched in tetraploids across a particular contrast were also likely to be among the list of significant GO terms in diploids (for the same contrast, Fig. 2e), in particular when comparing contrasts across watering conditions. These results suggest a relatively high degree of similarity in the functional response to abiotic stress, even if the specific set of genes undergoing a significant change in expression across abiotic conditions often differs considerably between cytotypes. The top-fifteen most significant GO categories in tetraploids, based on the list of differentially expressed genes in leaves when contrasting drought-stressed against well-watered plants at 28C (‘HD vs HW’; barplot in Figure 2e and Supplementary Materials 2 and 3), includes terms related to cell wall organization (GO:0071555) response to water deprivation (GO:0009414), mitotic cytokinesis (GO:0000281), as well as response to abscisic acid (GO:0009737) and auxin (GO:0009733), phytohormones known to play important roles in stress signaling and plant development (Zhao 2010, Osakabe et al. 2014, Kuromori et al. 2018). Most of these same categories were also enriched in 2X specimens (barplot in Fig. 2e).

There is evidence that plants activate distinct signaling and transcriptomic pathways when faced with different kinds of abiotic or biotic stresses, and that exposure to combinations of different stress factors activates unique regulatory circuits that cannot be extrapolated from the response to single stressors (reviewed in Suzuki et al. 2014). The results here presented for leaf tissue suggest that diploid and tetraploid specimens have a similar response during transition to drought at high temperature, a double-stress scenario, but diverge on their responses to single abiotic stress factors. Autopolyploidy is known to induce higher salt tolerance in *A. thaliana* (Chao et al. 2013), and higher drought resistance in *A. thaliana* (Del Pozo and Ramirez-Parra 2014, 2015), *Citrus limoni* (Allario et al. 2013), and *Hordeum marinum* (Zhou et al. 2019). *B. pubescens* and *B. pendula* are both susceptible to water stress (Atkinson 1992, Hemery et al. 2010) and while *B. pubescens* has been generally considered to be more sensitive to drought than *B. pendula,* this assertion has been recently challenged (González de Andrés et al. 2023). *B. pubescens* is better adapted to extremely low winter temperatures, being able to survive in Arctic regions, but *B. pendula* has a higher tolerance to high temperatures (Atkinson 1992, Hemery et al. 2010). It is unclear, however, whether current differences in abiotic stress responses between cytotypes are an outright consequence of genome duplication or, as inferred in some polyploid complexes (Maherali et al. 2009), have evolved through selection during the 1.6 My since the ancestors of these two birch species diverged (Leal e al. 2024).

### Transcriptomic Differences Between Diploids and Tetraploids Highlight Changes in Secondary Metabolism and Defense Response to Pathogens and Other Organisms

GO term enrichment analysis of genes differentially expressed between diploid and tetraploid individuals (Fig. 2b) identified several biological processes whose regulation has been remodeled in the aftermath of autopolyploidization (Fig. 3a and Supplementary Materials 4). There is a low to moderate degree of overlap between DEG associated to different abiotic conditions, between 170 and 216 genes, depending on tissue type (Figs. 3b-c), and only a relatively small number of GO terms is enriched across all contrasts (Fig. 3a). Across the eight abiotic and tissue-type scenarios studied, the most common highly enriched (lowest p-value) terms relate to the regulation of secondary metabolic processes (terms highlighted in blue in Fig. 3a), and defense/immune-system responses to pathogens and other organisms (terms highlighted in pink). Other highly enriched terms pertain to biological functions associated to recognition of pollen (GO:0048544) and heat acclimation (GO:0010286). These results are congruent with enrichment analysis performed in other plant diploid-autopolyploid complexes. Reprogramming of metabolic networks is among the most frequently functional changes observed in autopolyploids when comparing transcriptomic profiles across ploidies (Lu et al. 2006, Yu et al. 2010, Mu et al. 2012, Del Pozo and Ramirez-Parra 2014, Dai et al. 2015, Zhou et al. 2015, Braynen et al. 2017, Wang et al. 2018, Zhou et al. 2019). Expression variation in genes involved in biotic defense response is also a relatively common aftereffect of whole genome duplication, having been previously reported in *A. thaliana* (Yu et al. 2010, Del Pozo and Ramirez-Parra 2014), *Betula platyphylla* (Mu et al. 2012), *Morus alba* (Dai et al. 2015), and *Brassica rapa* (Braynen et al. 2017).

**Figure 3.**
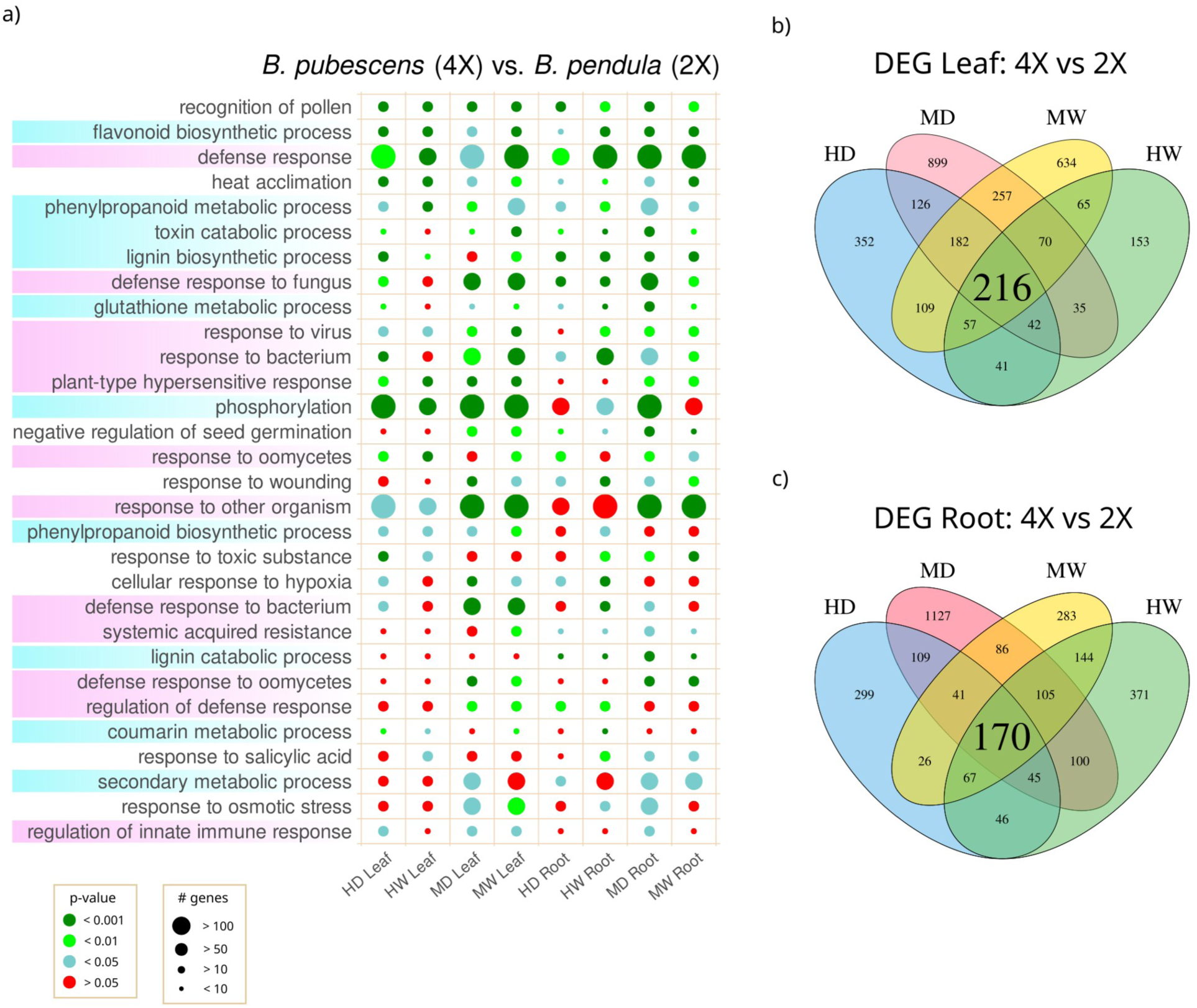
| Transcriptomic differences between diploids and tetraploids – (**a**) Top-30 most common enriched GO categories, based on set of differentially expressed genes (DEG) across ploidies, for different biotic conditions and tissue type. GO terms enriched in all contrasts are shown on top. GO terms associated to metabolic processes are highlighted in blue; terms associated to biotic defense response are highlighted in pink. Only GO categories containing at least 10 annotated genes were considered. See Supplementary Materials 4 for complete list. Number of genes refers to number of significant DEG in category. ‘D’, ‘W’, ‘H’, and ‘M’ stand for ‘drought’, ‘watered’, ‘hot(28C)’, and ‘mild(20C)’, respectively. (**b-c**) Venn diagrams showing the total number of DEG across ploidies, for different biotic conditions and tissue type. Only genes with p-adj < 0.05, |log2FoldChange| > 1, and baseMean > 10 were considered to be differentially expressed.

Changes in metabolic demands are expected to occur in polyploids as genome doubling invariably leads to changes in cell size, surface-to-volume ratios, number and size of organelles, cellular distances, cell division rates, and cell-cycle duration (reviewed in Doyle and Coate 2019). Production of alkaloids, terpenes and other secondary metabolites involved in chemical defense has been shown to increase dramatically in several medical plant autopolyploids (Levin 1983, Lavania et al. 2012, te Beest et al. 2012), providing a likely explanation for observations that whole genome duplication often confers greater resistance to pathogens and herbivores (Levin 1983, te Beest et al. 2012, Edger et al. 2015, van de Peer et al. 2021, Hagen and Mason 2024). Biochemical analysis of *B. pendula* and *B. pubescens* leaves revealed that *B. pubescens* produces specific classes of terpenes and flavonoids not found in diploid *B. pendula* (reviewed in Atkinson 1992) and that some flavonoids are found at much higher concentration in *B. pubescens* (Valkama et al. 2004), hinting at disparities in secondary metabolic profiles between the two species that might help explain differences in susceptibility to pathogenic fungi and insect herbivory (Valkama et al. 2005). Our results confirmed the presence of significant expression differences between cytotypes in genes involved in the flavonoid synthetic pathway (Fig. 3a), metabolism of one of its persecutors, phenylpropanoid, and identified one extra class of metabolites, glutathiones, known to play an important role in disease resistance in Arabidopsis (Parisy et al. 2007). The phenylpropanoid metabolic pathway is also involved in lignin biosynthesis, which is also enriched in our analysis (Fig. 3a). Lignin is a key component of the secondary cell wall and also acts as a defense polymer against microbial activity (Xie at al. 2018). Analysis of lignin content in cell walls performed in newly synthesized *A. thaliana* autopolyploid lines revealed that it correlates negatively with ploidy level (Corneillie et al. 2019), while studies of *A. thaliana* mutant lines show that reduced lignin content is often associated with upregulation of genes involved in pathogen resistance, likely due to cross-talk between lignin biosynthesis, growth, and defense pathways (reviewed in Xie at al. 2018).

In established natural autopolyploids such as *B. pubescens, A. arenosa*, and *Cardamine amara*, changes in biotic defense architecture are expected to occur as diploid and tetraploid cytotypes diverge and independently adapt to new environments. Such a development has been inferred in *C. amara*, where genomes scans comparing natural autotetraploid and diploid populations identified several genes involved in abiotic defense response located in genomic regions bearing a signature of selection (Bohutínská et al. 2021A). Interestingly, genome-wide DNA methylation analysis of diploid and newly synthesized *B. pendula* autotetraploids has revealed the presence of CHH hypermethylated regions in polyploids that are enriched for metabolic processes and biotic defense responses (Zhang et al. 2024), suggesting that some transcriptomic differences in genes involved in biotic defense occur instantly as a consequence of polyploidization.

It has been noted that WGD shares some common characteristics with endoreduplication (the programmed increase in ploidy level in somatic cells, Scholes and Paige 2015), even if their transcriptomic profiles are generally distinct (Robinson et al. 2018). Work carried out on *A. thaliana* mutant lines showed that induction of the endocycle activates metabolic pathways used in plant chemical defense (Mesa et al. 2017). The presence of commonalities was further confirmed in studies of endoreduplication in diploid and autotetraploid *A. arenosa* populations, which identified a small group of genes involved in both processes, and further revealed that DEG underlying endoreduplication are enriched for metabolic processes and defense responses (Wos et al. 2022). The extent to which polyploidization co-opts specific cell programs involved in endoreduplication, eg by monitoring total cellular DNA, remains poorly understood and merits further investigation.

### The Set of Meiosis Genes Showing Preferential Allelic Expression in *B. pubescens* Includes Several Genes Under Selection in *A. arenosa* and *A. lyrata* Autotetraploids

High-resolution allelic expression analysis of *B. pubescens* meiotic genes was performed with EAGLE-RC (Kuo et al. 2018), a method developed to determine the most likely origin of a read among a set of competing genomic hypothesis based on genotype differences, and which can be used to differentiate transcriptomic differences between alleles of distinct evolutionary origin (Kuo et al. 2020, Hu et al. 2021). The aim of this analysis was to determine whether expression in *B. pubescens* meiotic genes is strongly biased towards alleles introgressed from *B. humilis* or *B. nana*, as this would suggest that such genes could be under selection. For this purpose, reads were mapped to three different assemblies (*B. pendula*, *B. nana*, and *B. humilis*), and the EAGLE-RC pipeline was used to compute read likelihoods for each of the three reference assemblies before carrying out expression quantification. Reads mapping to the *B. pendula* assembly are expected to include those associated to alleles introgressed from *B. pendula* into *B. pubescens*, as well as reads that map to alleles descending from the ancestral *B. pubescens* population(s) at the time of speciation. The gene set under consideration includes homologues to all meiotic genes previously identified as being under selection in the autotetraploids *A. arenosa* (Yant et al. 2013, Wright et al. 2015, Monnahan et al. 2019, Bohutínská et al. 2021b), *A. lyrata* (Marburger et al. 2019), and *C. amara* (Bohutínská et al. 2021a), as well as homologues to 74 Arabidopsis meiotic genes based on lists compiled by Bohutínská et al. (2021a) and Mercier et al. (2015). We excluded meiotic genes not expressed in *B. pubescens* in the tissues and/or conditions covered in our experimental design, as well as genes for which homology with *A. thaliana* could not be established. In total, our analysis includes 59 meiotic genes, 11 of which are known to be under selection in one or both Arabidopsis autotetraploids (Supplementary Table S2).

Allelic expression analysis was first performed in diploid *B. pendula* individuals, where the vast majority of reads were expected to map to the *B. pendula* assembly. Our results confirm that for most meiotic genes expression is mainly detected in alleles of *B. pendula* origin, independently of tissue type, abiotic conditions, or ecotype (Fig. 4a and Supplementary Fig. S5a). Residual expression assigned to alleles of supposedly *B. nana* or *B. humilis* origin is in most cases likely due to assembly and mapping errors and/or lack of differentiation between assemblies. However, as there is evidence of low levels of introgression from other birch species into *B. pendula*, namely from *B. pubescens, B. nana*, and *B. platyphylla* (Palmé et al. 2004, Wang et al. 2014, Tsuda et al. 2017, Leal et al. 2024), it cannot be excluded that for some individuals, or even some genes, introgressed alleles have replaced one or both ancestral *B. pendula* alleles.

**Figure 4.**
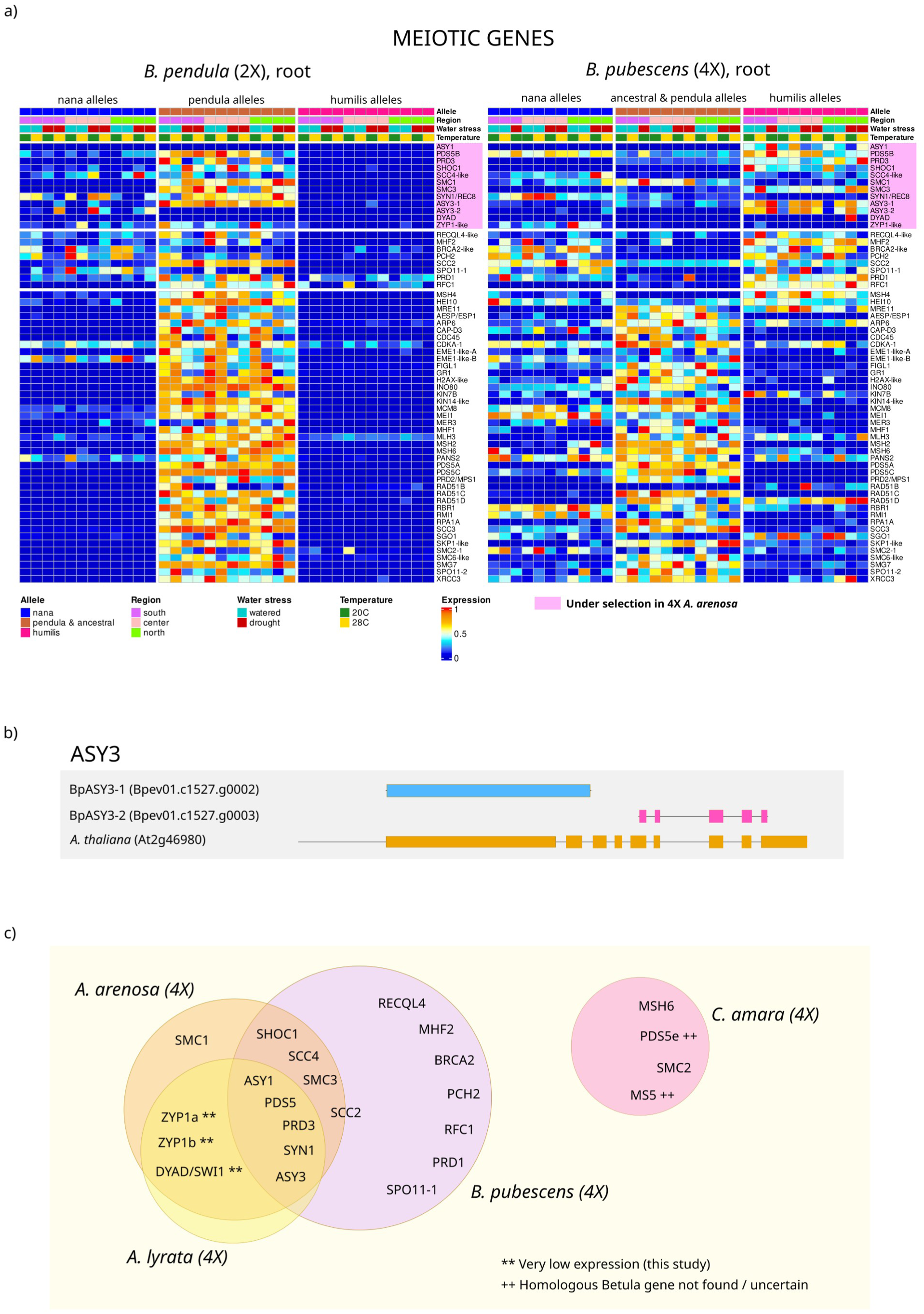
| Allelic expression in meiotic genes – (**a**) Heatmap of normalized expression levels for 59 meiotic genes (rows), assayed in *B. pendula* and *B. pubescens* root tissue, across different abiotic conditions, ecotypes (“Region”), and allelic evolutionary origin (“Allele”). Values for each cell are the average expression for three biological replicates. (**b**) Sequence alignment for *AtASY3*, *BpASY3-1*, and *BpASY3-2* performed with Clustal Omega based on exonic nucleotide sequences. (**c**) Venn diagram showing list of meiotic genes under selective pressure identified in *B. pubescens* (this study), and in *A. arenosa*, *A. lyrata*, and *C. amara* autotetraploids (see main text for references).

Expression analysis of *B. pubescens* specimens revealed an altogether different transcriptomic landscape where, for a large fraction of meiotic genes, alleles of *B. nana* or *B. humilis* origin were detected in at least some individuals (Fig. 4a and Supplementary Fig. S5b). This agrees with previous studies showing that although *B. pubescens* contains extensive genomic regions acquired from *B. nana* and *B. humilis*, most such introgression events occur stochastically and, with rare exceptions, most borrowed alleles are kept at low frequency (Leal et al. 2024).

The most striking result, however, is the presence of 16 *B. pubescens* meiotic genes where alleles introgressed from *B. nana* or, more prominently, *B. humilis*, were preferentially expressed, while alleles mapping to the *B. pendula* assembly were either not expressed or showed reduced expression (Fig 4a and Supplementary Fig. S5b). Absence of expression for specific allelic variants could be due to such alleles not being present, having been replaced by alleles originating from *B. humilis* or *B. nana*, or because expression is based on allele-inherent characteristics. Allele-specific expression has been previously observed in polyploids (Margarido et al. 2022) and diploid hybrids (Springer and Stupar 2007, Zhuang and Adams 2007, Hu et al. 2016), and is attributed to differential regulation of alleles of different origin, likely due to selection. Be that as it may, for these 16 genes, biased expression favoring alleles borrowed from *B. humilis* or *B. nana* was generally observed, independently of abiotic conditions and ecotype, strongly suggesting that the genomic regions where these genes are located are under strong selective pressure. Of these 16 meiotic loci, five (*ASY1*, *PDS5B*, *PRD3*, *SYN1*, and *ASY3*) were previously shown to exhibit strong signatures of a selective sweep following WGD in both 4X *A. arenosa* and 4X *A. lyrata* (Yant et al. 2013, Wright et al. 2015, Marburger et al. 2019, Monnahan et al. 2019), with three others (*SMC3, SHOC1, SCC4*) having been identified as also being under selection in *A. arenosa* autotetraploids (Yant et al. 2013, Bohutínská et al. 2021b). One meiotic gene identified here as having biased allelic expression, *SCC2*, has been suggested to be under selection in *A. arenosa* autopolyploids based on genomic scans contrasting 4X *A. arenosa* against diploid *A. lyrata* and *A. thaliana* populations (Hollister et al. 2012). As this gene has not been flagged in any subsequent study carried out in Arabidopsis autotetraploids, which all contrasted autopolyploids against their diploid progenitors, this would suggest that *SCC2* is likely under selection also in diploid *A. arenosa*. *DYAD* and *ZYP1*, two meiotic genes under selection in 4X *A. arenosa* and 4X *A. lyrata*, showed only vestigial levels of expression in the tissues and conditions included in this study (Fig. 4a and Supplementary Fig. S5a).

A particularly intriguing case concerns *ASY3*, a gene implicated in synaptonemal complex formation between homologous chromosomes. A specific haplotype of this core meiotic gene has been revealed to be widely shared between *A. lyrata* and *A. arenosa* autotetraploids, with the former believed to have provided the latter with a pre-adapted variant via introgressive hybridization (Seear et al. 2020). Together with *ASY1*, this is also one of the few meiotic genes for which there is direct evidence of a link between natural allelic variation and meiotic stability in Arabidopsis autotetraploids, with specific *ASY3* allelic variants shown to induce a reduction in meiotic crossover rates during metaphase I (Morgan et al. 2020, Seear et al. 2020). Low crossover rates lead to the suppression of multivalent formation, a chromosomal pairing configuration often associated with genetic disorders such as aneuploidy, and the reestablishment of (random) bivalent pairing in the autopolyploid (Bomblies et al. 2016). In birch, *ASY3* has been replaced by two separate genes (here named as *BpASY3-1* and *BpASY3-2*) which together cover most of the nucleotide sequence span of their *A. thaliana* homologue (Fig. 4b). Expression of the ancestral/pendula allele is generally null or subsided for both *BpASY3* genes

Other *B. pubescens* meiotic genes shown here to have biased allelic expression favoring alleles of *B. humilis* or *B. nana* origin include *RECQL4*, *MHF2*, *BRCA2*, *PCH2*, *RFC1*, *PRD1*, and *SPO11-1* (Fig. 4a and Supplementary Fig. S5b). *RECQL4* and *MHF2* are anti-crossover factors, *RFC1* and *PCH2* modulate class I crossover formation (also class II for *PCH2*), *SPO11-1* and *PRD1* are required for double strand break formation during meiotic recombination, and *BRCA2* is involved in double strand break repair (Wang et al. 2012, Lambing et al. 2015, Mercier et al. 2015). It is possible that some of these genes are not themselves under selection but are located near loci under strong selective pressure, an issue that warrants further investigation. In general, the set of meiotic genes listed above are believed to be located in different *B. pubescens* chromosomes or are otherwise placed far apart when located in the same chromosome (Supplementary Table S3). The exceptions are the two *ASY3* genes, which are located adjacent to each other, and *SMC3* and *RFC1,* which are located only about 200k bp apart. Allelic expression bias is identical for *SMC3* and *RFC1*, showing a preference for the allele of *B. humilis* origin (Fig. 4a and Supplementary Table S5, discussed below).

Allelic mapping as carried out here is performed concurrently against three birch assemblies based on individuals sampled from a natural population. As such, the assemblies used could themselves contain loci introgressed from other birch species, which would have biased our analysis. Furthermore, borrowed alleles found in *B. pubescens* might be relatively rare, or have been lost, in the diploid species of origin. We consequently carried out an additional set of analyses where the assemblies used during mapping were based on different accessions (Supplementary Tables S4). While transcriptomic patterns varied slightly based on the specific accessions used during mapping, the same set of 16 meiotic genes showing preferential expression towards alleles of *B. humilis* or *B. nana* origin was generally flagged (Supplementary Tables S5). Several other meiotic genes showing biased allelic expression were occasionally detected. For the most part, such instances were sporadic and seemed to reflect unique genomic characteristics of the specific accessions used as a reference during mapping. More importantly, overall these results suggest that, among *B. pubescens* meiotic genes showing biased expression, 9 genes preferentially express borrowed alleles of *B. humilis* origin, a set that includes *ASY1*, *PDS5B*, *PRD3*, *SHOC1*, *SMC3*, *RECQL4*, *MHF2*, *PCH2*, and *RFC1* (Supplementary Tables S5). Only one meiotic gene, *SYN1*, showed strong preference for expressing alleles introgressed from *B. nana*. Three meiotic genes (*SCC4*, *SCC2*, and *SPO11-1*) favored expression of both alleles, while no clear trend could be discerned for the remain three meiotic genes (*ASY3*, *BRCA2*, *PRD1*).

## Conclusions

In this article, we provide evidence of a conserved evolutionary response to whole genome duplication in angiosperms, confirming genomic shifts, both at the functional and gene level, first identified in the Arabidopsis model system. This was done by analyzing phenotypic and transcriptomic differences between autotetraploid *B. pubescens* and its closest diploid relative, *B. pendula*, two birch species 120-140 Mya divergent from the *Arabidopsis* genus. Specifically, we identified 16 meiotic genes in *B. pubescens* whose expression is strongly biased towards alleles introgressed from *B. humilis* or *B. nana*, suggesting that they could be under selective pressure. Among these, there were 5 genes (*ASY1*, *PDS5*, *PRD3*, *SYN1*, and *ASY3*) previously shown to be under selection in 4X *A. arenosa* and 4X *A. lyrata* (Fig. 4c), and four others (SMC3, *SHOC1*, *SCC2*, and *SCC4*) believed to be under selection in autotetraploid *A. arenosa*. The overlap between the set of meiotic genes putatively under selection in *B. pubescens* and in the two Arabidopsis autotetraploid species is striking, in particular if we take into account that three of the four remaining meiotic Arabidopsis genes (*ZYP1a*, *ZYP1b*, and *DYAD/SWI1*) (Yant et al. 2013, Wright et al. 2015, Marburger et al. 2019, Monnahan et al. 2019, Bohutínská et al. 2021b) could not be evaluated in the presented study as they were not expressed (or were otherwise only very weakly expressed) in the tissues/conditions included in the experimental design (Fig. 4c). There was no overlap with the set of meiotic genes detected in 4X *C. amara* (Fig. 4c), which could be due to weak or noisy selection signals in the genome scans carried out in *C. amara* or, as speculated by the authors, because, unlike in Arabidopsis autotetraploids, 4X *C. amara* might have been provided with a set of pre-adapted meiotic genes by its diploid progenitor (Bohutínská et al. 2021a). The exact functional role played by the meiotic genes here discussed remains to be elucidated for the most part, both in Arabidopsis and in other autopolyploid plant species, an endeavor that is certain to further our understanding of the evolutionary forces and constrains faced by flowering plants in the aftermath of genome duplication.

Further confirming their affinity with other autopolyploid-diploid complexes, the two birch species here studied showed identical reactions to abiotic stress in terms of stomatal activity, photosynthesis rates, and leaf ABA concentration, and their transcriptomes were for the most part undifferentiated. Cross-ploidy differences in gene expression were relatively limited and generally dominated by functional categories associated to cell secondary metabolism and resistance to pathogens. These results are in broad agreement with those observed in other autopolyploid plant species (Warner and Edwards 1993, Yu et al. 2010, Mu et al. 2012, Allario et al. 2013, Del Pozo and Ramirez-Parra 2014, Dai et al. 2015, Braynen et al. 2017, Wos et al. 2019), and are congruent with a model where changes in cell size triggered by genome duplication lead to stoichiometric adjustments in metabolic processes affecting cell-wall composition and, through cross-talk between (cell-wall) polymer biosynthesis and chemical defense pathways, the reconfiguration of biotic defense responses. This posits changes in defense architecture as one of the key fundamental challenges, together with increased meiotic instability, that must be overcome by most, if not all, neo-autopolyploids if they are to survive and leave progeny. Taken together, these results give support to the view that the path to successful autopolyploidization is highly conserved in flowering plant, as hypothesized by Bohutínská et al. (2021a), and that it may often depend on the ability of a nascent autopolyploid to obtain specific allelic variants from non-parental species via introgression, as previously shown in Arabidopsis by Marburger et al. (2019) and Seear et al. (2020).

## Materials and Methods

### Plant material and growth conditions

Autotetraploid *B. pubescens* and diploid *B. pendula* seeds collected from different geographic regions (Fig. 1a) were grown in a programmable LTCB-19 growth chamber (BioChambers) under controlled conditions of 23 h light (150 µmol m^−2^ s^−1^) and 1 h dark photoperiod, at 21°C daytime and 18 °C nights. Seeds were cold-stratified for one month in the dark at 5 °C before being sown in SW Horto Yrkesplantjord soil and transferred to a growth chamber for germination. After six months, plants were subjected to a five-month vernalization cycle (T_min_ = 6 °C). After bud burst, specimens were kept for five weeks under pre-vernalization temperate and light conditions before being randomly divided into four groups and subjected to different abiotic stress scenarios: (i) MW: 20°C/watered; (ii) HW: 28°C/watered; (iii) MD: 20°C/drought; (iv) HD: 28°C/drought (Fig. 1b). For each of these four treatments, each species is represented by nine specimens (3 ecotypes, with 3 biological replicates per ecotype), adding up to a grand total of 72 individuals. Light conditions were kept unchanged (23h light; 1h dark). Soil moisture was monitored daily using a ProCheck (Decagon Devices) hand-held sensor, with watering levels being adjusted for each individual specimen in order to maintain well-watered plants at 50-60% VWC (volumetric water content) and soil in plants under drought at 5-20% VWC. Individual plants were shifted within growth chambers daily in order to reduced local light and temperature effects. After being subjected to abiotic stress for five weeks, leaf and root tissue used in mRNA and phytohormone quantification were collected from each of the 72 specimens 15-18 hours after lights-on, promptly frozen in liquid nitrogen, and kept in storage at -80 °C. Leaf temperature was measured 24h prior to sample collection using a IR270 Infrared Thermometer (Extech Instruments).

### RNA extraction, library preparation and sequencing

Total RNA was extracted from leaf and root samples using a method combining a CTAB lysis buffer and the Spectrum Total Plant RNA extraction kit (Sigma), following the protocol described in Carrell et al. (2022). Sequencing libraries were prepared from 450ng total RNA using a TruSeq Stranded mRNA Kit (Illumina) including polyA selection and unique dual indexes, according to the manufacturer’s protocol (#1000000040498). Libraries were sequenced on paired-end mode (150 bp) on an Illumina NovaSeq 6000 platform (Illumina). Details about each accession can be found in Supplementary Materials 5.

### Additional WGS accessions

Multiple WGS accessions were used to generate the three reference assemblies (*B. pendula*, *B. humilis*, and *B. nana*) required when carrying out allele-specific expression analysis (described below). Some of these accessions were part of a previous study (Salojärvi et al. 2017). For the new WGS *B. nana* and *B. pendula* accessions, total DNA was extracted from bud and leaf tissues using a DNeasy Plant Mini Kit (Qiagen AG), according to the instructions provided by the manufacturer apart from the incubation during lysis which was extended to 2 hours. Individual libraries were sequenced on paired-end mode (150 bp) on an Illumina NovaSeq 6000 platform (Illumina). For the *B. humilis* accessions, genomic DNA was extracted using a DNAsecure Plant Kit (Tiangen Biotech) according to the manufacturer’s instructions and paired-end short reads (150 bp) were generated for each sample on an Illumina HiSeq X Ten platform (Illumina). Further details about WGS accessions used can be found in Supplementary Table S4.

### Quality filtering, read mapping, and gene expression quantification

Initial filtering of RNA-Seq raw reads was performed with TrimGalore v0.6.1 (Krueger 2015), a wrapper tool around CUTADAPT (Martin 2011), with the intent of removing adapter sequences and trimming of low-quality read ends. PloyA tails were removed using PRINSEQ v0.20.4 (Schmieder and Edwards 2011), and SortMeRNA v2.1b (Kopylova et al. 2012) was used to identify and filter out reads produced by residual rRNA fragments. Reads shorter than 30 bp after trimming were filtered out. Read mapping was carried out with STAR v2.7.2b (Dobin et al. 2013) in two-pass mode and using default settings. *B. pubescens* and *B. pendula* reads were mapped separately. Reads were mapped to the *B. pendula* reference assembly, vBpev01 (Salojärvi et al. 2017), using the associated annotation file. The average number of reads per sample decreased from 45.6 million (raw libraries) to 39.7 million after filtering, while the average read length decreased from 151 to 136 bp. For *B. pubescens* samples, at least 92.5% (mean = 95.5%) of reads mapped to the reference genome, and on average 92.3% of reads mapped to unique genomic positions (for *B. pendula* libraries, these values were 94.1%, 96.6% and 94%, respectively). Transcriptome assembly and gene expression quantification was performed with StringTie v1.3.3 (Pertea et al. 2015) in expression estimation mode and using the *B. pendula* annotation file to guide the assembly process. The final (un-normalized) gene count matrix, which reports the total number of sequencing reads that map to each gene for each sample, was generated with the prepDE.py script included in the StringTie suite.

### Differential gene expression analysis

Identification of differentially expressed genes across contrasts was performed with the R/Bioconductor package DESeq2 v1.42.1 (Love et al. 2014), based on the gene count matrix generated using StringTie. P-values were adjusted for multiple testing using the Benjamini-Hochberg correction with an alpha < 0.05 false discovery rate. Only genes with p-adj < 0.05, |log2FoldChange| > 1, and baseMean > 10 were considered to be differentially expressed. Depending on the contrast, between three and nine biological replicates were used per condition (specific values provided in main text). Genes with very low coverage (mean count across all samples ≤ 1) and genes for which expression was observed in just one single sample were filtered out before carrying out the analysis. The DESeq2 design formula used to estimated dispersions and fold changes included a correction term for “ecotype” when comparing expression levels across abiotic conditions or ploidy. A pairwise comparison model was used when evaluating expression levels across tissue type, as leaf and root tissues were collected from the same set of individuals. Normalization factors for each sample (size factors), which correct for library size, were estimated using the median ratio method as implemented in the default DESeq function, which assumes that only a fraction of all genes are deferentially expressed (Love et al. 2014). When used to compare expression levels in samples obtained from individuals of different ploidy, this approach allows for detection of changes in transcript stoichiometry (relative change) while avoiding some of the pitfalls associated to some well-known alternative library normalization procedures (Coate and Doyle 2015, Visger et al. 2019). In the approach here employed, it is assumed that the number of transcripts is generally proportional to gene copy number, with genes being flagged as differentially expressed if expression ratios across ploidies deviate significantly from this rule. Additionally, it was also assumed that transcript lengths remain unchanged across ploidies, a reasonable assumption given their relatively recent divergence (Leal et al. 2024). Principal component analysis (PCA) of the normalized count matrix was performed with the function plotPCA included with the DESeq2 package based on the top 2,000 genes with the highest variance.

### K-means cluster analysis

When producing heatmaps displaying normalized expression values across contrasts, K-means cluster analysis was used to partition the set of differentially expressed genes in groups with similar expression patterns across experimental conditions. Clustering analysis was carried out using the k-means algorithm as implemented on MATLAB, v. 2019a (Mathworks), following the procedure described in Leal et al. (2024).

### Gene ontology (GO) enrichment analysis

Identification of biological functions enriched across treatments was performed with the R/Bioconductor package TOPGO, v. 2.48.0 (Alexa et al. 2006), based on the set of differential expressed genes identified using DESeq2. GO annotation of *B. pendula* genes was based on the annotation of homologous genes in the *A. thaliana* model species. Identification of protein homologs between the two species was carried out with HMMER v3.4 (Eddy 2011). GO functional annotation of *A. thaliana* genes was obtained from the Uniprot database (https://www.uniprot.org/uniprotkb?query=%28taxonomy_id%3A3701%29, downloaded January 25, 2024). Statistical significance of overrepresented GO categories was estimated using the elim Fisher’s exact test (Fisher-elim) implemented in topGO and should be interpreted as corrected for multiple testing (Alexa and Rahnenfuhrer 2024).

### Allele-specific expression analysis

Allelic expression analysis of *B. pubescens* and *B. pendula* genes was performed with EAGLE-RC v1.1.1 (Kuo et al. 2018). For each library, reads were independently mapped to three different reference assemblies (*B. pendula*, *B. humilis*, and *B. nana*) using the STAR mapper, and the EAGLE-RC pipeline was subsequently used to compute and compare read likelihoods for each assembly in order to determine the read origin. The *B. humilis* and *B. nana* reference assemblies used during read mapping were based on consensus sequences generated on the basis of individual WGS accessions (Supplementary Table S4) by applying all SNP and indel variants particular to each accession to the *B. pendula* reference genome, following the procedure detailed in Leal et al. (2023). The *B. pendula* reference assembly used during allelic mapping was either the official *B. pendula* reference genome or was otherwise based on consensus sequences generated in a similar fashion to those representing *B. humilis* and *B. nana*.

Read quantification was performed with featureCounts, included in the Subread v2.0.3 package (Liao et al. 2013), producing three count matrixes, one for each reference assembly. When applied to *B. pubescens* samples, two of the count matrices quantify expression by introgressed alleles of *B. humilis* and *B. nana* origin, respectively. The third count matrix includes read counts associated to alleles introgressed from *B. pendula*, as well as to *B. pubescens* ancestral alleles. The latter are expected to map best to the *B. pendula* reference assembly (among the three assemblies included) as *B. pendula* is one of its two closest extant relatives (Leal et al. 2024), the other being *B. platyphylla*, found in East Asia. Normalization of each of the count matrices produced by Subread was carried out by first adding all three count matrices, estimating sample size factors for each library based on the cumulative matrix using the function estimateSizeFactors in DESeq2, and then applying these size factors to all three individual count matrices.

### Chromosomal assignment and sequence alignment of meiotic loci

Chromosomal assignment of meiotic loci was performed on the basis of the *B. pendula* chromosomal map, v5834 (Salojärvi et al., forthcoming). Alignment of *A. thaliana* and *B. pendula ASY3* exonic nucleotide sequences was performed with Clustal Omega (Sievers and Higgins 2021). Exonic nucleotide sequences for *AtASY3* (At2g46980) were downloaded from the Arabidopsis TAIR database (https://www.arabidopsis.org/locus?key=31711, last accessed 2024-06-25).

### Ploidy determination

The ploidy level of each *B. pubescens* and *B. pendula* seedling was initially verified using a CytoFLEX flow cytometer (Beckman Coulter), with leaf samples prepared with the CyStain UV Precise P kit (Sysmex) following the instructions provided by the manufacturer. Ploidy levels were verified again after RNA sequencing by checking the distribution of read counts at biallelic variant sites, as described in Yoshida et al. (2013) and Leal et at. (2024). Identification of SNP variants based on RNA-Seq data was performed following the GATK RNA-Seq short variant discovery workflow (Brouard and Bissonnette 2022, GATK Team 2024). As *B. pendula* can be found in the 2X and 4X forms in nature, we also confirmed that all tetraploid individuals included in this study were *B. pubescens* by checking their genomic composition using genomic polarization (Leal et al. 2023), a phasing technique, as described in Leal et al. (2024). Unlike 4X *B. pendula*, *B. pubescens* contains sizable genomic components introgressed from *B. humilis* and *B. nana* that can be quantified using genomic polarization (Supplementary Materials 6).

### Leaf gas exchange measurements

Net photosynthetic rate (A), leaf respiration rate (E), and stomatal conductance (g_s_) were measured on canopy leaves using a LCpro-T portable infrared gas analyzer (ADC BioScientific) at a CO_2_ concentration (C_set_) of 400 ppm under light saturated conditions, Q_set_ = 500 μmol m^-2^ s^-1^. The temperature in the leaf chamber (T_set_) was set to either 20°C or 28°C, depending on the specimen being measured. Water-vapor pressure into the chamber (e_ref_) was 15.0±0.6 mBar (20°C, drought), 17.6±0.8 mBar (28°C, drought), 15.9±0.5 mBar (20°C, watered), and 18.1±0.7 mBar (28°C, watered). Before performing readings, rates of CO_2_ exchange were allowed to equilibrate until steady-state conditions were attained (usually 3 to 5 minutes). Measurements were performed 5 weeks after inception of water and temperature treatment (10 weeks after bud break), 15-18 hours after lights-on.

### Measurement of leaf ABA content

Concentration of abscisic acid (ABA) in leaf samples was determined by Ultra-High-Performance Liquid Chromatography coupled with Triple Quadrupole Mass Spectrometry (UHPLC-QqQ-MS). For further details see Supporting Materials 1. Leaf samples were deep-frozen in liquid nitrogen immediately upon collection and kept at -80C until phytohormone extraction and analysis. Sample collection took place 5 weeks after inception of water and temperature treatment, 15 hours after lights-on.

### Statistical analysis of phenotypic data

Differences in stomatal function, photosynthesis, and ABA concentration across ploidy and abiotic conditions were evaluated by performing multi-way analysis of variance (ANOVA) tests with the R-package emmeans v1.9.0 (Lenth 2024) based on the best linear model selected using the AIC criteria. P-values were adjusted for multiple comparisons using the Tukey’s HSD Test.

### Data Accessibility

RNA-Seq raw sequence reads from this study have been uploaded to the European Bioinformatics Institute (EMBL-EBI) data-base ArrayExpress (https://www.ebi.ac.uk/arrayexpress/) under accession number XXXXXXX [data is being held private until publication]. WGS raw sequence reads are available at the The European Nucleotide Archive (ENA) at www.ebi.ac.uk and can be accessed with the XXXXXXX [data is being held private until publication] and PRJEB14544 accession codes. Supplementary scripts and data are available from the Dryad Digital Repository XXXXXXX and the GitHub repository https://github.com/LLN273/Allelic.Expression_4X_vs_2X.

## Acknowledgements

We are very grateful to Dana Carper and Sara Jawdy for providing the RNA extraction protocol. We also thank Tianlin Duan, Marion Orsucci, Erik Gudmunds, and Javier Florenza García for their help and suggestions during lab work, Nathalie Zeballos for assistance during seed sampling, and Yanara Marincevic-Zuniga for coordinating the RNA sequencing workflow. The Swedish Metabolomics Centre, Umeå, Sweden (www.swedishmetabolomicscentre.se) is acknowledged for quantification of plant hormones by LC-QqQ-MS. RNA Sequencing was performed by the SNP&SEQ Technology Platform in Uppsala, part of the National Genomics Infrastructure (NGI) Sweden and Science for Life Laboratory. DNA Sequencing was performed at Novogene. Some of the genome sequences used in this article were generated within the project ‘Tree for ME’ which is financed by the Swedish Energy Agency. The computations were enabled by resources in project NAISS 2024/22-11 provided by the National Academic Infrastructure for Supercomputing in Sweden (NAISS) at UPPMAX, funded by the Swedish Research Council through grant agreement no. 2022-06725.

## Author contributions

J.L.L, M.L. and J.S conceived of the study. J.L.L. and E.H. carried out seed collection with support from M.L. J.L.L. performed growth chamber experiments, phenotypic measurements, and tissue sampling with assistance and advice from G.G., A.B., D.M.E., and P.M. J.L.L. performed RNA extraction and characterization with assistance and advice from A.B. and D.M.E. M.P.H.E. and J.C. performed DNA extraction. J.L.L. performed all data analyses. Paper was written by J.L.L. with input from all authors. The final version of the manuscript was approved by all co-authors

