## Supplementary_Materials_1 for "Conserved Evolutionary Response to Whole Genome Duplication in Angiosperms Revealed Using High Resolution Gene Expression Profiling"

**Measurement of leaf ABA content [Detailed description]**

Chemicals

Solvents: Methanol, HPLC-grade was obtained from Fischer Scientific (Waltham, MA, USA) Acetonitrile and 2-Propanol hypergrade for LC-MS LiChrosolv was obtained from Merck (Darmstadt, Germany), Acetic acid (glacial), Aldrich Chemical Company, Inc. (Milwaukee, WI, USA), H2O Milli-Q. Abscisic Acid (ABA) hormone standards were acquired from (Olchemim.cz, Olomouc, Czech Republic). Internal standards Abscisic Acid-D6 (ABA-D6) was acquired from Cayman Chemicals (Ann Arbor, USA). A 9 level calibration of curve ABA was prepared by serial dilution (ranging: 62.5 fg/µl - 400 pg/µl) and spiked with internal standards (0.1 ng/µl).

Sample Preparation

For extraction 1 ml extraction mix (80% MeOH, 1% Acetic acid) was added to approximately 15 mg sample material. The sample was shaken with one tungsten beads at 30 Hz for 3 min in a mixer mill (MM 400, Retsch) whereupon the beads were removed. Samples were centrifuged at 14 000 rpm (18 620g) for 10 min at +4 °C and the supernatant was transferred to vials and stored at -80°C. A small aliquot of the remaining supernatants were pooled and used to create quality control (QC) samples. Blank samples, i.e. samples without starting material, were prepared the same way as the plant samples. The extracted samples was purified on Oasis HLB cartridges (1 cc, 30 mg, Waters, Milford, MA, USA). Prior to SPE purification 600 or 700µL of the extract (based on the amount of starting material) was spiked with 50µL internal standard mix (ABA-D6 at 0.04ng/µL), evaporated in a vacuum concentrator (miVac Quattrol Concentrator, Genevac™) until approximately 80µL remained, before loading on the HLB cartridges the samples were spiked with 5µL 1M HCl. The SPE purification was performed at a Pressure + 48 system from Biotage (Uppsala, Sweden) at 2 - 5 psi positive pressure. The HLB columns were equilibrated with 1 ml 10% MeOH before loading the acidified samples. After washing with 2 ml 10%MeOH, the samples were eluted with 2 x 1 ml 80% MeOH + 1% Acetic acid. After elution the samples were evaporated until dryness in a vacuum concentrator and stored at -80°C until analysis.

Analysis by UHPLC-QQQ-MS

The analysis was conducted using an Agilent UHPLC system (Infinity 1290) coupled with an electrospray ionization source (ESI) to an Agilent 6495 triple quadrupole system equipped with iFunnel Technology (Agilent Technologies, Santa Clara, CA, USA). Chromatographic separation was performed on a Waters UPLC HSS T3 column (2.1 mm × 50 mm, 1.8-µm particle size), the mobile phase consisted of 0.1% acetic acid in MQ-water (A) and ACN (B). The gradient was 15% B for 5 min followed by a linear gradient from 15 to 50% over 2.5 minutes, then increased from 50 to 99% B in one minute, hold at 99% B from 8.5-9.5 min, and thereafter returned to initial conditions and re-equilibrated for 2 minutes. The flow rate was set of 600 µL/min and the column was heated to 40 °C. Prior to analysis the samples were dissolved in 40 µl 40 % MeOH, the injection volume was 10 µL. The mass spectrometer was operated in positive/ negative switching ESI mode with gas temperature set at 150°C; gas flow 12 L min-1; nebulizer pressure 20 psi; sheath gas temperature 400°C; sheath gas flow 12 L min-1; capillary voltage 4000 V (neg), 3500 V (pos); nozzle voltage 1500V (pos), 500 V (neg); iFunnel high pressure RF 150 V; iFunnel low pressure RF 60 V. The fragmentor voltage 380 V and cell acceleration voltage 5 V. Data were processed using MassHunter Qualitative Analysis and Quantitative Analysis (QqQ; Agilent Technologies, Atlanta, GA, USA) and Excel (Microsoft, Redmond, Washington, USA) software. Different extract volumes and mg of starting material were compensated for during calculations.

**Table S1** | Sampling locations

| Chromosome No. | Species | Country | Latitude | Longitude | Group | Climate[a] |
|---|---|---|---|---|---|---|
| 2n=2X=28 | *B. pendula* | Sweden | 56.02 | 13.88 | South | Temperate oceanic |
| | *B. pendula* | Sweden | 58.41 | 15.1 | South | Temperate oceanic |
| | *B. pendula* | Sweden | 62.16 | 17.35 | Center | Climatic contact zone |
| | *B. pendula* | Sweden | 63.13 | 18.4 | Center | Climatic contact zone |
| | *B. pendula* | Sweden | 65.5 | 21.22 | North | Subarctic |
| | *B. pendula* | Sweden | 66.31 | 20.57 | North | Subarctic |
| 2n=4X=56 | *B. pubescens* | Sweden | 56.02 | 13.88 | South | Temperate oceanic |
| | *B. pubescens* | Sweden | 56.75 | 14.55 | South | Temperate oceanic |
| | *B. pubescens* | Sweden | 58.41 | 15.1 | South | Temperate oceanic |
| | *B. pubescens* | Sweden | 64.53 | 21.07 | Center | Subarctic |
| | *B. pubescens* | Sweden | 65.5 | 21.22 | Center | Subarctic |
| | *B. pubescens* | Norway | 69.11 | 23.25 | North | Subarctic (above arctic circle) |

[a] Köppen-Geiger climate classification

**Table S2 |** List of meiotic genes included in allelic expression analysis. Homology between *A. thaliana* and *B. pendula* genes established using HMMER (see *Materials and Methods*).

| Protein name (*A. thaliana*) | *A. thaliana* Uniprot entry | *B. pendula* gene name | *A. thaliana* length (aa) | *B. pendula* length (aa) | E-value (hmmsearch) |
|---|---|---|---|---|---|
| ASY1 | F4HRV8 | Bpev01.c0052.g0112 | 596 | 594 | 3.50E-255 |
| PDS5B | F4I735 | Bpev01.c0340.g0016 | 1,410 | 1,397 | 0 |
| PRD3 | Q0WWX5 | Bpev01.c1519.g0005 | 449 | 381 | 8.00E-84 |
| SHOC1 | F4KG50 | Bpev01.c1342.g0004 | 1,594 | 1,647 | 0 |
| SCC4-like | Q9FGN7 | Bpev01.c0089.g0042 | 726 | 505 | 6.50E-231 |
| SMC1 | Q6Q1P4 | Bpev01.c0050.g0118 | 1,218 | 1,753 | 0 |
| SMC3 | Q56YN8 | Bpev01.c0429.g0017 | 1,204 | 1,204 | 0 |
| SYN1/REC8 | Q9S7T7 | Bpev01.c0115.g0080 | 627 | 600 | 3.80E-156 |
| ASY3 | Q0WR66 | Bpev01.c1527.g0002<br>Bpev01.c1527.g0003 | 793 | 599<br>153 | 1.70E-47<br>3.60E-27 |
| DYAD | Q9FGN8 | Bpev01.c0477.g0006 | 639 | 446 | 7.40E-128 |
| ZYP1-like | Q9LME2 | Bpev01.c0151.g0010 | 871 | 661 | 3.40E-206 |
| RECQL4-like | Q8L840 | Bpev01.c0320.g0008 | 1,188 | 1,322 | 0 |
| MHF2 | Q8L7N3 | Bpev01.c0349.g0010 | 104 | 104 | 2.50E-44 |
| BRCA2-like | Q7Y1C4 | Bpev01.c0154.g0060 | 1,155 | 1,110 | 0 |
| PCH2 | Q8H1F9 | Bpev01.c0126.g0062 | 467 | 463 | 1.50E-240 |
| SCC2 | A5HEI1 | Bpev01.c0135.g0027 | 1,846 | 1,835 | 0 |
| SPO11-1 | Q9M4A2 | Bpev01.c0112.g0012 | 362 | 469 | 3.80E-142 |
| PRD1 | O23277 | Bpev01.c0016.g0159 | 1,330 | 1,309 | 0 |
| RFC1 | Q9C587 | Bpev01.c0429.g0032 | 956 | 1,008 | 0 |
| MSH4 | F4JP48 | Bpev01.c0939.g0006 | 792 | 743 | 0 |
| HEI10 | F4HRI2 | Bpev01.c1420.g0005 | 304 | 294 | 7.30E-169 |
| MRE11 | Q9XGM2 | Bpev01.c1138.g0010 | 720 | 723 | 0 |
| AESP/ESP1 | Q5IBC5 | Bpev01.c0190.g0016 | 2,180 | 2,210 | 0 |
| ARP6 | Q8LGE3 | Bpev01.c0126.g0003 | 421 | 437 | 3.00E-213 |
| CAP-D3 | O24610 | Bpev01.c1260.g0014 | 1,314 | 1,158 | 0 |
| CDC45 | Q9LSG6 | Bpev01.c0645.g0005 | 596 | 662 | 0 |
| CDKA;1 | P24100 | Bpev01.c0957.g0013 | 294 | 294 | 1.80E-181 |
| EME1-like-A | C5H8J1 | Bpev01.c1024.g0001 | 551 | 380 | 8.20E-82 |
| EME1-like-B | C5H8J1 | Bpev01.c1024.g0002 | 551 | 385 | 1.30E-92 |
| FIGL1 | F4JEX5 | Bpev01.c0566.g0059 | 680 | 682 | 1.30E-301 |
| GR1 | Q9ZRT1 | Bpev01.c1456.g0009 | 588 | 641 | 2.80E-129 |
| H2AX-like | Q9S9K7 | Bpev01.c0804.g0006 | 142 | 144 | 5.20E-88 |
| INO80 | Q8RXS6 | Bpev01.c0029.g0149 | 1,507 | 1,537 | 0 |
| KIN7B | Q8LNZ2 | Bpev01.c0051.g0186 | 938 | 949 | 0 |
| KIN14-like | Q9LX99 | Bpev01.c0286.g0029 | 1,273 | 1,284 | 0 |
| MCM8 | Q9SF37 | Bpev01.c0288.g0020 | 801 | 746 | 0 |
| MEI1 | A0A1P8AWF3 | Bpev01.c0464.g0026 | 1,001 | 994 | 0 |
| MER3 | Q5D892 | Bpev01.c0062.g0040 | 1,133 | 1,231 | 0 |
| MHF1 | Q9FI55 | Bpev01.c1202.g0026 | 242 | 140 | 3.20E-59 |
| MLH3 | F4JN26 | Bpev01.c0022.g0150 | 1,155 | 1,158 | 3.40E-255 |
| MSH2 | O24617 | Bpev01.c2021.g0001 | 937 | 1,024 | 0 |
| MSH6 | O04716 | Bpev01.c0129.g0106 | 1,324 | 1,324 | 0 |
| PANS2 | Q94CK6 | Bpev01.c0223.g0031 | 194 | 188 | 4.50E-30 |
| PDS5A | B6EUB3 | Bpev01.c2470.g0002 | 1,605 | 1,753 | 0 |
| PDS5C | Q8GUP3 | Bpev01.c1406.g0006 | 873 | 679 | 4.60E-148 |
| PRD2/MPS1 | F4KDF5 | Bpev01.c0602.g0003 | 377 | 372 | 4.20E-138 |
| RAD51B | Q9SK02 | Bpev01.c0448.g0003 | 370 | 372 | 8.00E-191 |
| RAD51C | Q8GXF0 | Bpev01.c0192.g0013 | 363 | 346 | 9.50E-182 |
| RAD51D | Q9LQQ2 | Bpev01.c0396.g0007 | 322 | 321 | 1.40E-127 |
| RBR1 | Q9LKZ3 | Bpev01.c0457.g0045 | 1,013 | 1,018 | 0 |
| RMI1 | Q5XUX6 | Bpev01.c0557.g0037 | 644 | 602 | 4.10E-176 |
| RPA1A | Q9SKI4 | Bpev01.c0051.g0101 | 640 | 621 | 0 |
| SCC3 | O82265 | Bpev01.c0863.g0002 | 1,098 | 1,125 | 0 |
| SGO1 | F4J3S1 | Bpev01.c0191.g0036 | 572 | 477 | 2.90E-30 |
| SKP1-like | Q39255 | Bpev01.c0022.g0074 | 160 | 155 | 6.70E-91 |
| SMC2-1 | Q9C5Y4 | Bpev01.c0668.g0023 | 1,175 | 1,176 | 0 |
| SMC6-like | Q9FLR5 | Bpev01.c0668.g0019 | 1,058 | 1,058 | 0 |
| SMG7 | A9QM73 | Bpev01.c0223.g0049 | 1,059 | 987 | 0 |
| SPO11-2 | Q9M4A1 | Bpev01.c0170.g0013 | 383 | 382 | 4.80E-205 |
| XRCC3 | Q9FKM5 | Bpev01.c0094.g0093 | 304 | 296 | 6.10E-124 |

**Table S3 |** Genomic position of *B. pubescens* genes with biased allelic expression, based on *B. pendula* chromosomal map.

| Protein name | Bp gene | Chromosome | start | stop |
|---|---|---|---|---|
| ASY1 | Bpev01.c0052.g0112 | Chr1 | 1,418,408 | 1,423,842 |
| PDS5B | Bpev01.c0340.g0016 | Chr1 | 35,712,182 | 35,740,222 |
| PRD3 | Bpev01.c1519.g0005 | Chr9 | 21,012,632 | 21,019,383 |
| SHOC1 | Bpev01.c1342.g0004 | Chr3 | 14,885,459 | 14,895,752 |
| SCC4 | Bpev01.c0089.g0042 | Chr12 | 24,849,566 | 24,859,503 |
| SMC1 | Bpev01.c0050.g0118 | Chr13 | 1,427,635 | 1,461,938 |
| SMC3 | Bpev01.c0429.g0017 | Chr12 | 4,547,281 | 4,592,416 |
| SYN1/REC8 | Bpev01.c0115.g0080 | Chr5 | 922,824 | 929,484 |
| ASY3-1 | Bpev01.c1527.g0002 | Chr8 | 27,037,782 | 27,039,581 |
| ASY3-2 | Bpev01.c1527.g0003 | Chr8 | 27,041,290 | 27,044,147 |
| RECQ4A | Bpev01.c0320.g0008 | Chr10 | 17,689,670 | 17,713,010 |
| MHF2 | Bpev01.c0349.g0010 | Chr9 | 23,700,405 | 23,701,834 |
| BRCA2B | Bpev01.c0154.g0060 | Chr11 | 1,733,742 | 1,740,902 |
| PCH2 | Bpev01.c0126.g0062 | Chr9 | 22,953,878 | 22,961,279 |
| SCC2 | Bpev01.c0135.g0027 | Chr3 | 5,373,351 | 5,391,670 |
| SPO11-1 | Bpev01.c0112.g0012 | Chr9 | 1,260,278 | 1,278,856 |
| PRD1 | Bpev01.c0016.g0159 | Chr1 | 8,659,244 | 8,666,466 |
| RFC1 | Bpev01.c0429.g0032 | Chr12 | 4,824,696 | 4,857,306 |

**Table S4** | Sample name, species, sampling location, ENA accession, and mean sequence coverage of extra accessions used to generate the three reference assemblies employed when performing allele-specific expression analysis with EAGLE-RC.

| Sample ID | Species | Country | Latitude | Longitude | ENA Run Accession | Mean sequencing coverage | Notes |
|---|---|---|---|---|---|---|---|
| Bh1489 | *B. humilis* Schrk. | Russia | 53.75 | 119.77 | | 20 | natural population |
| Bh1493 | *B. humilis* Schrk. | Russia | 53.75 | 119.77 | | 22 | natural population |
| Bh1500 | *B. humilis* Schrk. | Russia | 53.75 | 119.77 | | 23 | natural population |
| Bh1503 | *B. humilis* Schrk. | Russia | 53.75 | 119.77 | | 19 | natural population |
| A002-2* | *B. nana* L. | Finland | | | ERR2026268 | 24 | Natural Resources Institute Finland |
| ERR179410** | *B. nana* L. | Scotland | 57.17 | -4.77 | ERR179410 | 25 | |
| NANAS9501 | *B. nana* L. | Sweden | 57.47 | 12.67 | | 24 | natural population |
| NANAS9504 | *B. nana* L. | Sweden | 57.47 | 12.67 | | 22 | natural population |
| NANAS9506 | *B. nana* L. | Sweden | 57.47 | 12.67 | | 25 | natural population |
| DJU212 | *B. pendula* Roth. | Sweden | 56.02 | 13.88 | | 31 | natural population |
| Reference genome* | *B. pendula* Roth. | | | | | | *B. pendula* reference genome vBpev01 |
| Loimaa1* | *B. pendula* Roth. | Finland | 61.42 | 23.10 | ERR2026257 | 39 | natural population |

* Part of a previous study (Salojärvi et al. 2017).
* Part of a previous study (ENA ERR179410).

**Table S5 |** *B. pubescens* genes with biased allelic expression, as identified using the EAGLE-RC pipeline, for different reference assembly combinations.

| EAGLE-RC mapping (accessions) | | | Gene name | | | | | | | | | |
|---|---|---|---|---|---|---|---|---|---|---|---|---|
| ***B. pendula*** | ***B. nana*** | ***B. humilis*** | ASY1 | PDS5B | PRD3 | SHOC1 | SCC4 | SMC1 | SMC3 | SYN1 | ASY3-1 | ASY3-2 |
| DJU212 | ERR179410 | 1503 | hum | nh | hum | hum | nana | | hum | nana | hum | hum |
| DJU212 | ERR179410 | 1489 | hum | nana | | hum | nana | | hum | nana | hum | nh |
| DJU212 | A002-2 | 1503 | hum | hum | hum | hum | nh | | | nana | nana | hum |
| DJU212 | A002-2 | 1493 | hum | hum | hum | hum | nh | | | nana | nana | hum |
| RefGen | ERR179410 | 1500 | hum | nh | hum | hum | nh | | hum | nana | hum | hum |
| RefGen | ERR179410 | 1489 | hum | nana | nh | hum | nh | | hum | nana | hum | nh |
| RefGen | A002-2 | 1503 | hum | hum | hum | hum | nh | | hum | nana | nana | hum |
| RefGen | A002-2 | 1489 | hum | hum | hum | hum | nh | | hum | nana | nana | hum |
| RefGen | NAN9506 | 1489 | hum | hum | hum | hum | nh | | hum | nana | nh | nana |
| RefGen | NAN9501 | 1489 | hum | hum | nana | hum | nh | | nh | nana | nana | nh |
| RefGen | NAN9504 | 1489 | hum | hum | nana | hum | | | hum | nana | nana | nana |
| Loimaa1 | ERR179410 | 1489 | hum | nana | | hum | nana | | hum | nana | hum | nh |
| Loimaa1 | A002-2 | 1500 | hum | hum | | hum | nh | | | nana | nana | hum |

humilis allele over-expressed
nana allele over-expressed
nana & humilis alleles over-expressed

| EAGLE-RC mapping (accessions) | | | Gene name | | | | | | | |
|---|---|---|---|---|---|---|---|---|---|---|
| ***B. pendula*** | ***B. nana*** | ***B. humilis*** | RECQL4 | MHF2 | BRCA2 | PCH2 | SCC2 | SPO11-1 | PRD1 | RFC1 |
| DJU212 | ERR179410 | 1503 | hum | hum | nh | nh | nana | nh | hum | hum |
| DJU212 | ERR179410 | 1489 | hum | hum | nana | hum | nana | nh | nh | hum |
| DJU212 | A002-2 | 1503 | hum | hum | hum | hum | nh | nh | nh | hum |
| DJU212 | A002-2 | 1493 | hum | hum | hum | hum | nh | nh | | hum |
| RefGen | ERR179410 | 1500 | hum | hum | nh | hum | nh | nh | nh | hum |
| RefGen | ERR179410 | 1489 | hum | hum | nana | hum | nh | nh | nh | hum |
| RefGen | A002-2 | 1503 | hum | hum | hum | hum | nh | nh | hum | hum |
| RefGen | A002-2 | 1489 | hum | hum | hum | hum | nh | nh | nana | hum |
| RefGen | NAN9506 | 1489 | hum | nana | nh | hum | nh | nh | nana | nh |
| RefGen | NAN9501 | 1489 | hum | nana | nana | hum | nh | nh | nana | nh |
| RefGen | NAN9504 | 1489 | hum | nana | nh | hum | nh | nh | nana | hum |
| Loimaa1 | ERR179410 | 1489 | hum | hum | nana | hum | | nh | | hum |
| Loimaa1 | A002-2 | 1500 | hum | hum | hum | hum | nh | nh | | hum |

| EAGLE-RC mapping (accessions) | | | Gene name | | | | | | | | | |
|---|---|---|---|---|---|---|---|---|---|---|---|---|
| ***B. pendula*** | ***B. nana*** | ***B. humilis*** | MSH4 | HEI10 | MRE11 | ARP6 | CAP-D3 | CDKA-1 | MCM8 | EME1-A | EME1-B | MEI1 |
| DJU212 | ERR179410 | 1503 | hum | | | | | | | | | nana |
| DJU212 | ERR179410 | 1489 | hum | | | | | | | | | nana |
| DJU212 | A002-2 | 1503 | | nana | | | | | | | | |
| DJU212 | A002-2 | 1493 | | | | | | | | | | |
| RefGen | ERR179410 | 1500 | hum | | hum | nana | | | | nana | nana | nana |
| RefGen | ERR179410 | 1489 | hum | | hum | nana | | | | | | nana |
| RefGen | A002-2 | 1503 | | nana | nana | nana | | | | | | |
| RefGen | A002-2 | 1489 | | nana | nh | nana | | | | | | hum |
| RefGen | NAN9506 | 1489 | nh | nana | | nana | nana | hum | nana | | | nh |
| RefGen | NAN9501 | 1489 | hum | hum | | nana | nana | | nana | | | nana |
| RefGen | NAN9504 | 1489 | nh | hum | | | nana | | | | | hum |
| Loimaa1 | ERR179410 | 1489 | hum | | | | | nana | | | | |
| Loimaa1 | A002-2 | 1500 | | nana | | | | nana | | | | |

| EAGLE-RC mapping (accessions) | | | Gene name | | | | | | | | | | |
|---|---|---|---|---|---|---|---|---|---|---|---|---|---|
| ***B. pendula*** | ***B. nana*** | ***B. humilis*** | MER3 | MLH3 | MSH6 | PANS2 | PDR2 | PDS5C | RAD51B | RMI1 | RPA1A | SCC3 | SGO1 |
| DJU212 | ERR179410 | 1503 | | | | | | | | | | | |
| DJU212 | ERR179410 | 1489 | | | | | | | | | | | |
| DJU212 | A002-2 | 1503 | | | | | hum | | | | | | |
| DJU212 | A002-2 | 1493 | | | nana | | | | | | | | |
| RefGen | ERR179410 | 1500 | hum | | | nana | | | nana | nana | | | hum |
| RefGen | ERR179410 | 1489 | | hum | | nana | | | nana | | | | hum |
| RefGen | A002-2 | 1503 | hum | | | | hum | | | | | nana | nana |
| RefGen | A002-2 | 1489 | hum | | | | | | | hum | | nana | |
| RefGen | NAN9506 | 1489 | hum | nana | nana | nana | | nana | | | nana | | |
| RefGen | NAN9501 | 1489 | | | | | hum | nana | | | | | |
| RefGen | NAN9504 | 1489 | | | | | | | | | | | hum |
| Loimaa1 | ERR179410 | 1489 | nana | nana | | | | | | | | | hum |
| Loimaa1 | A002-2 | 1500 | hum | | | | | | | | | | |

**Figure S1** | Measures of (a) stomatal conductance; (b) transpiration rate; (c) and net photosynthetic rate, measured in *B. pendula* (2X) and *B. pubescens* (4X), across different ecotypes (south, center, north) and abiotic stress conditions [temperature (20C or 28C); watering (watered (WET) or drought]. Bar high indicates geometric mean for three replicates; error bars show 95% confidence interval of the mean.

a)

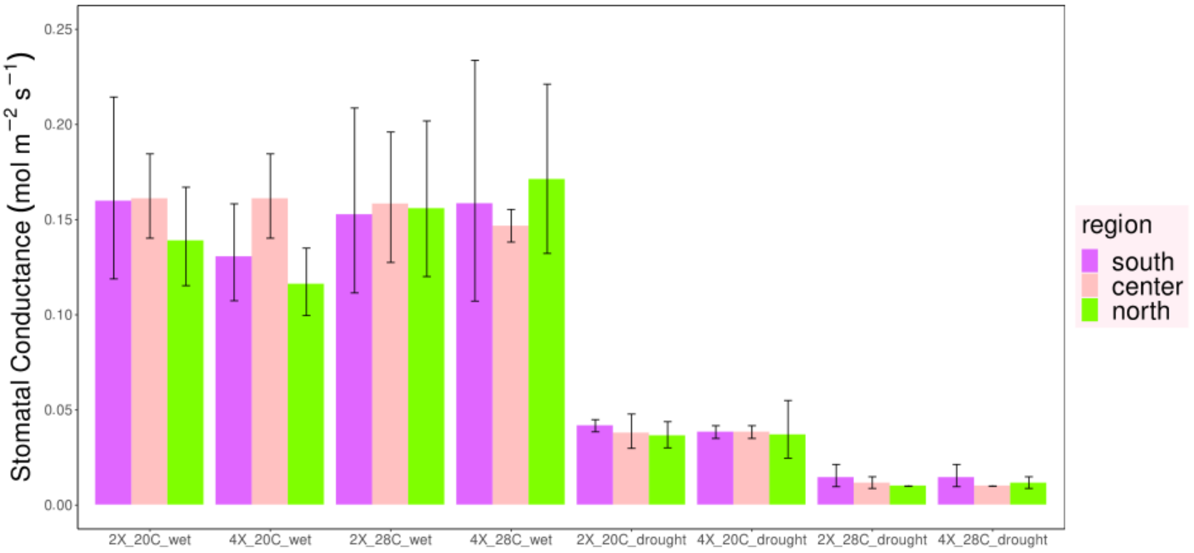


b)

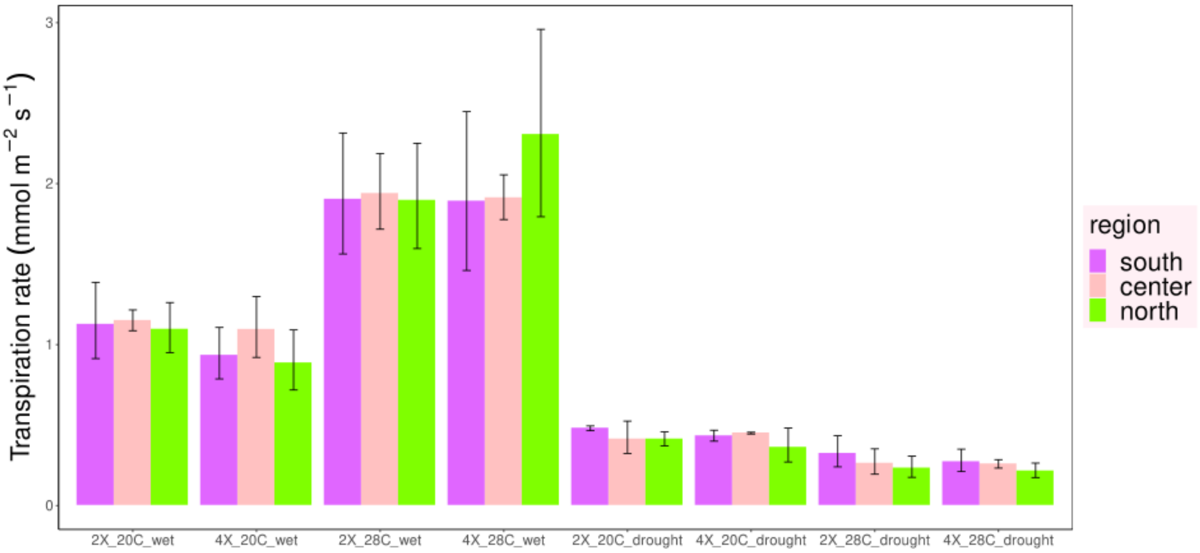

c)

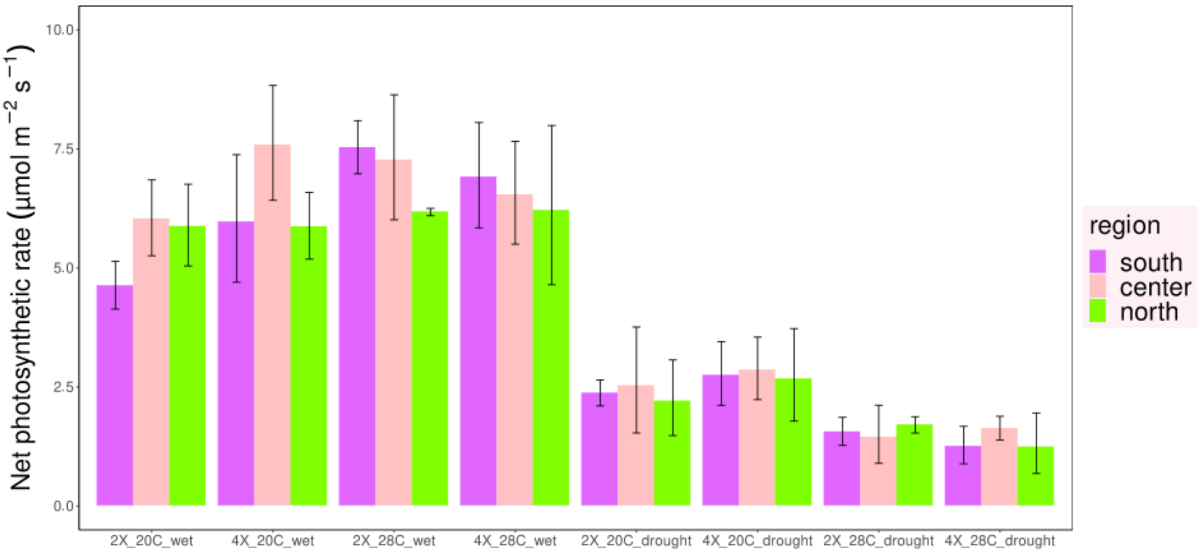

Net photosynthetic rate (µmol m−2 s−1)
10.0
7.5
5.0
2.5
0.0
2X_20C_wet
4X_20C_wet
2X_28C_wet
4X_28C_wet
2X_20C_drought
4X_20C_drought
2X_28C_drought
4X_28C_drought
region
south
center
north

**Figure S2 |** Leaf temperature in (a) lower and (b) upper canopy measured under different abiotic conditions. Bar high indicates geometric mean for nine replicates; error bars show 95% confidence interval of the mean. Symbols above bars show results from of multi-way ANOVA significance test based on the best linear model, corrected for multiple pairwise comparisons using the Tukey's HSD test. Parameter significance values (best linear model): '***': p-value < 0.001; '**': p-value < 0.01; '*': p-value < 0.05; 'ns': p-value > 0.05.

a)

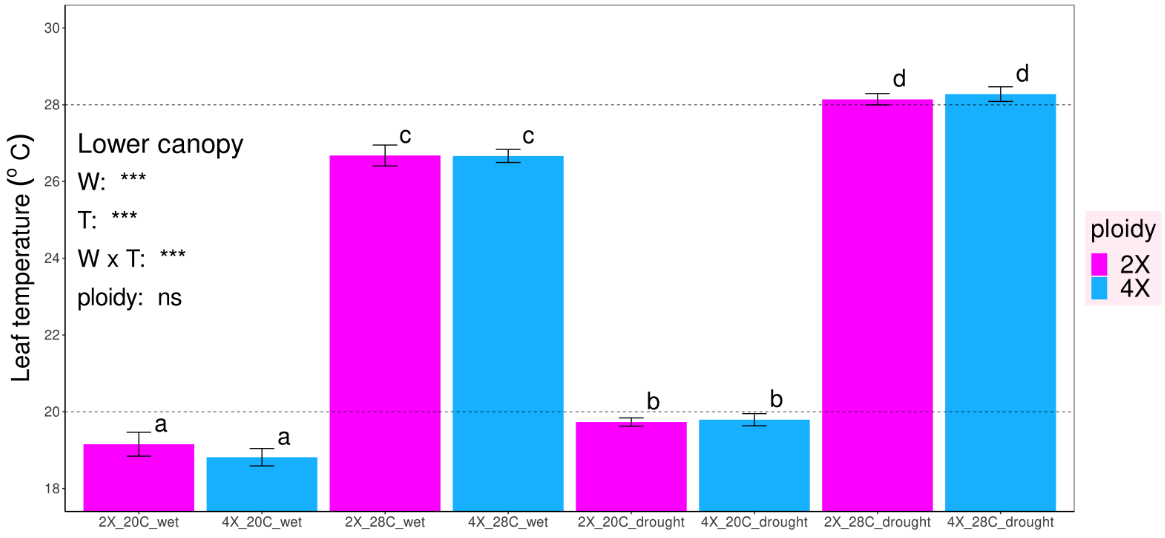


b)

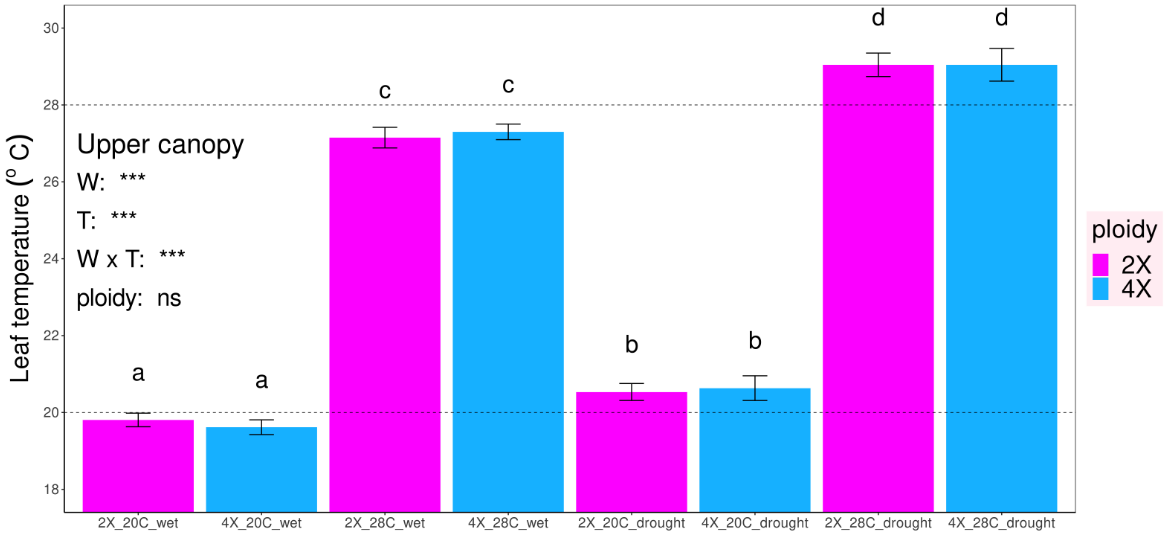

**Figure S3** | Heatmap of normalized expression levels for 16,890 differentially expressed genes (rows) for diploid and tetraploid samples, across different plant tissues and abiotic conditions (columns). Global expression patterns are dominated by differences across tissues (leaf vs root), followed by watering conditions (watered vs drought). Values for each cell are the average expression for nine biological replicates, apart from *B. pubescens* root samples subjected to drought at T=28C (seven replicates). Only genes with p-adj < 0.05, |log2FoldChange| > 1, and baseMean > 10, observed in at least one of the contrasts, were considered to be differentially expressed.

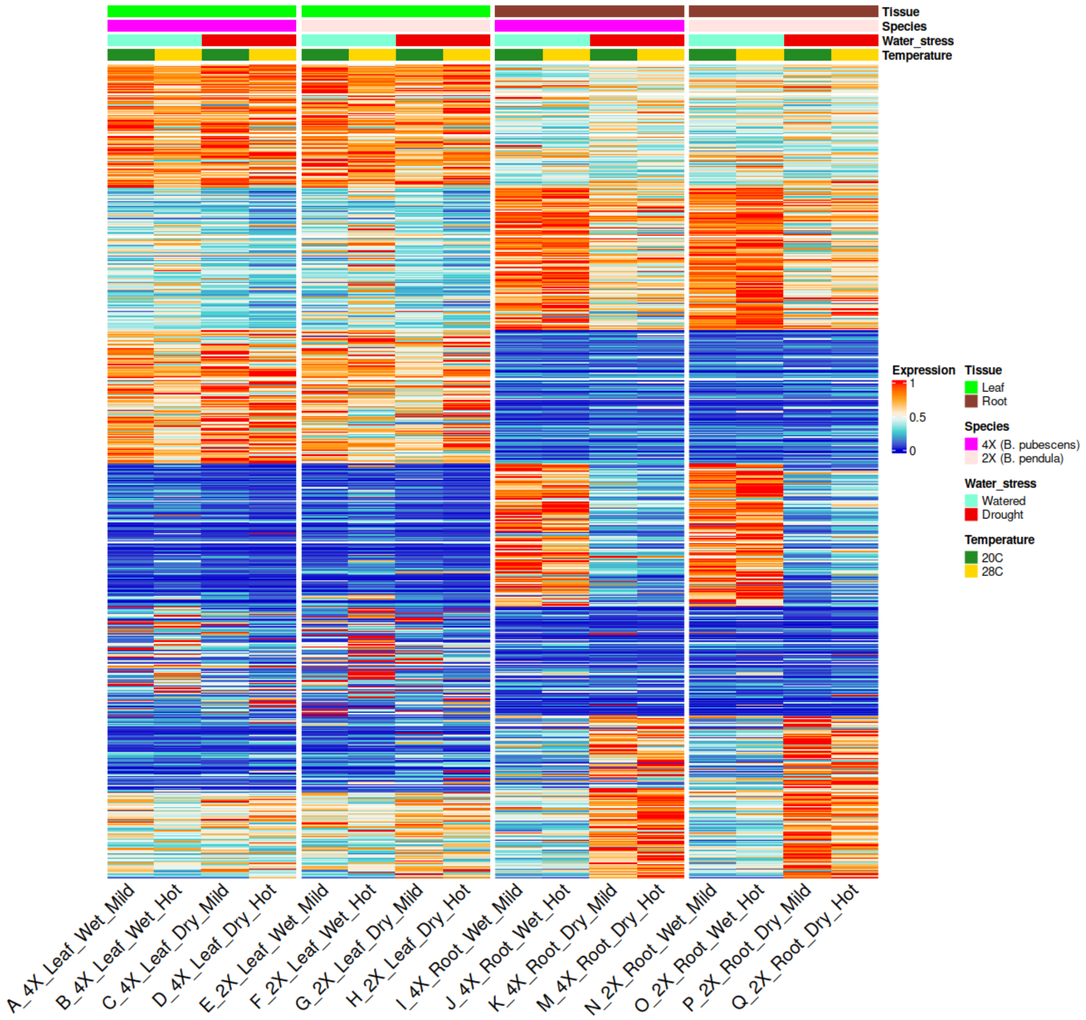

**Figure S4 |** Number of differential expressed genes across different ploidies (*B. pubescens* (4X); *B. pendula* (2X)), abiotic conditions, and plant tissue. Only genes with p-adj < 0.05, |log2FoldChange| > 1, and baseMean > 10 were considered to be differentially expressed.

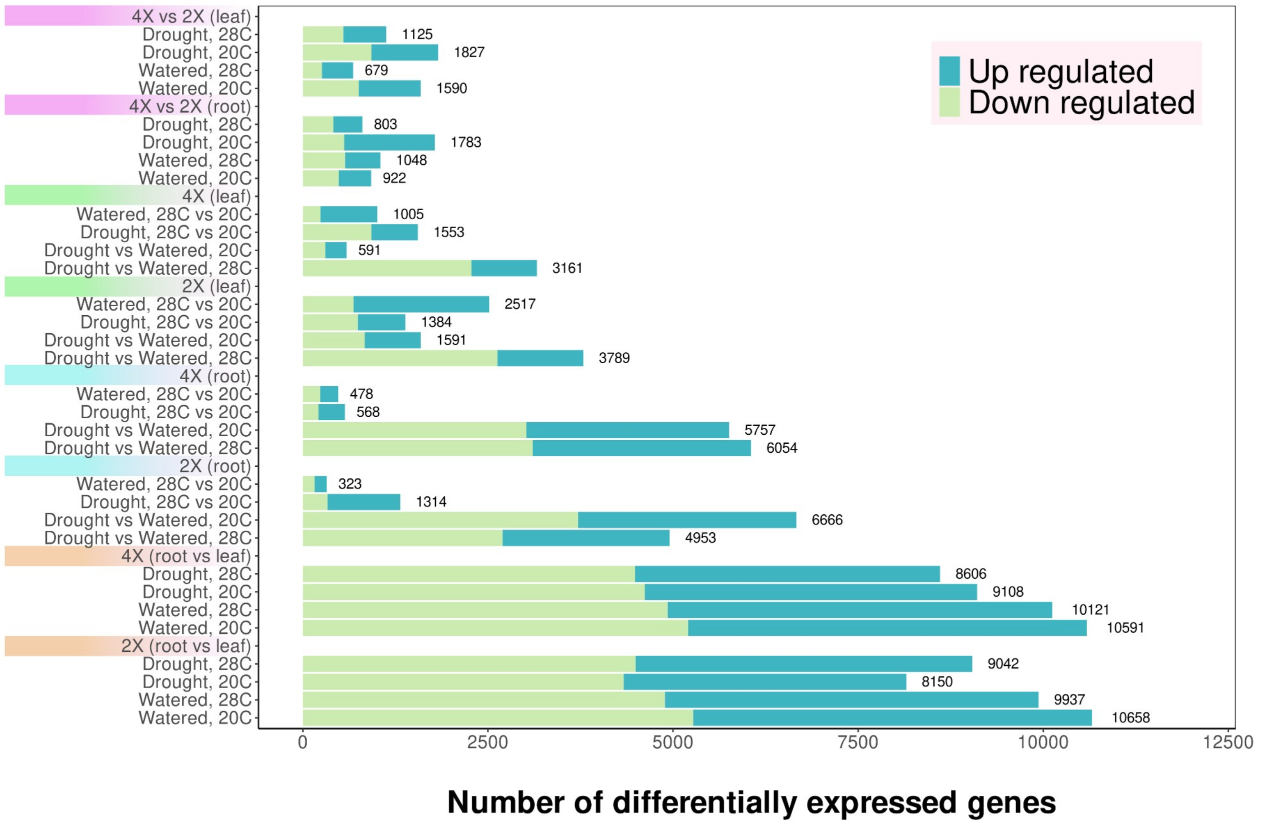

**Figure S5** | Heatmap of normalized expression levels for 59 meiotic genes (rows), assayed in (**a**) *B. pendula* and (**b**) *B. pubescens* leaf tissue, across different abiotic conditions, ecotypes ("Region"), and allelic evolutionary origin ("Allele"). Values for each cell are the average expression for three biological replicates.

(a)

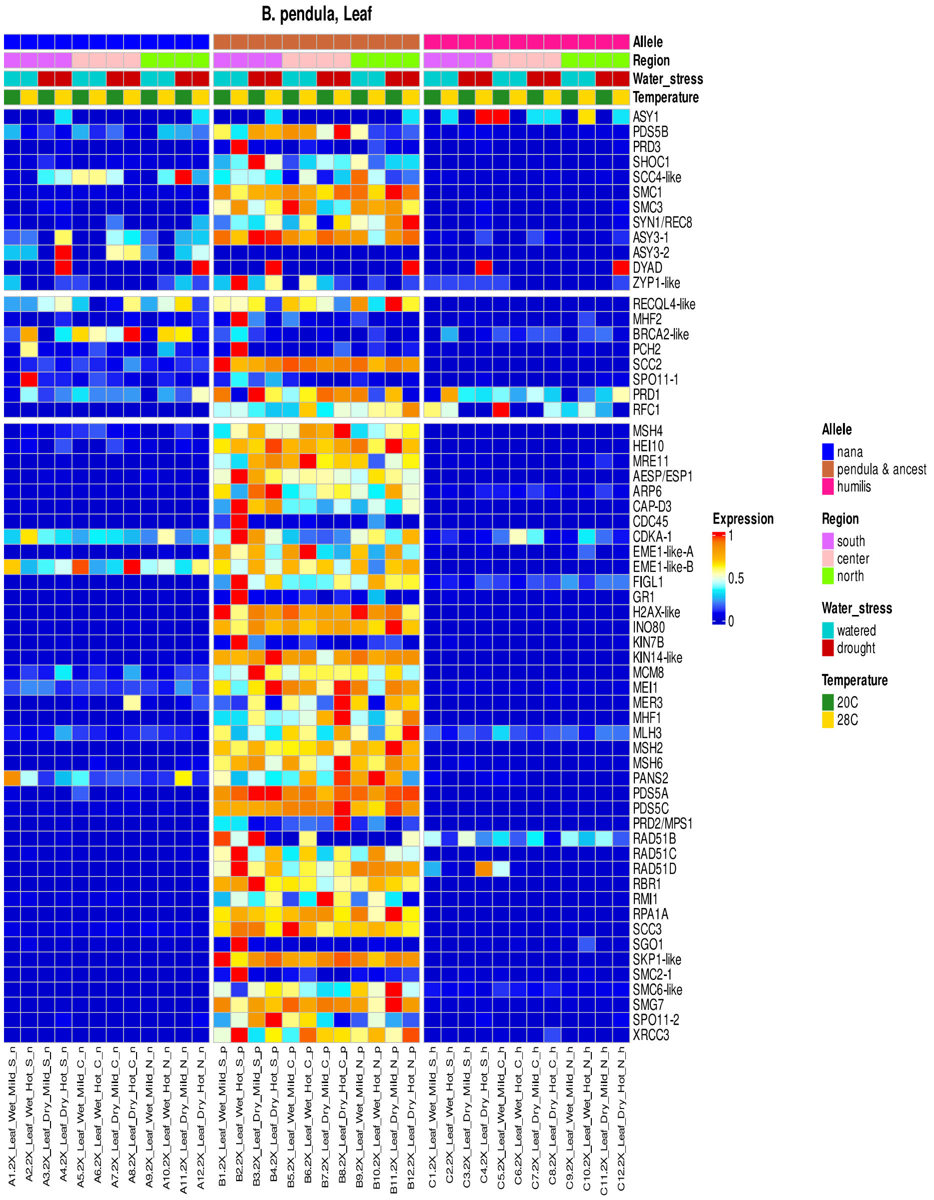

(b)

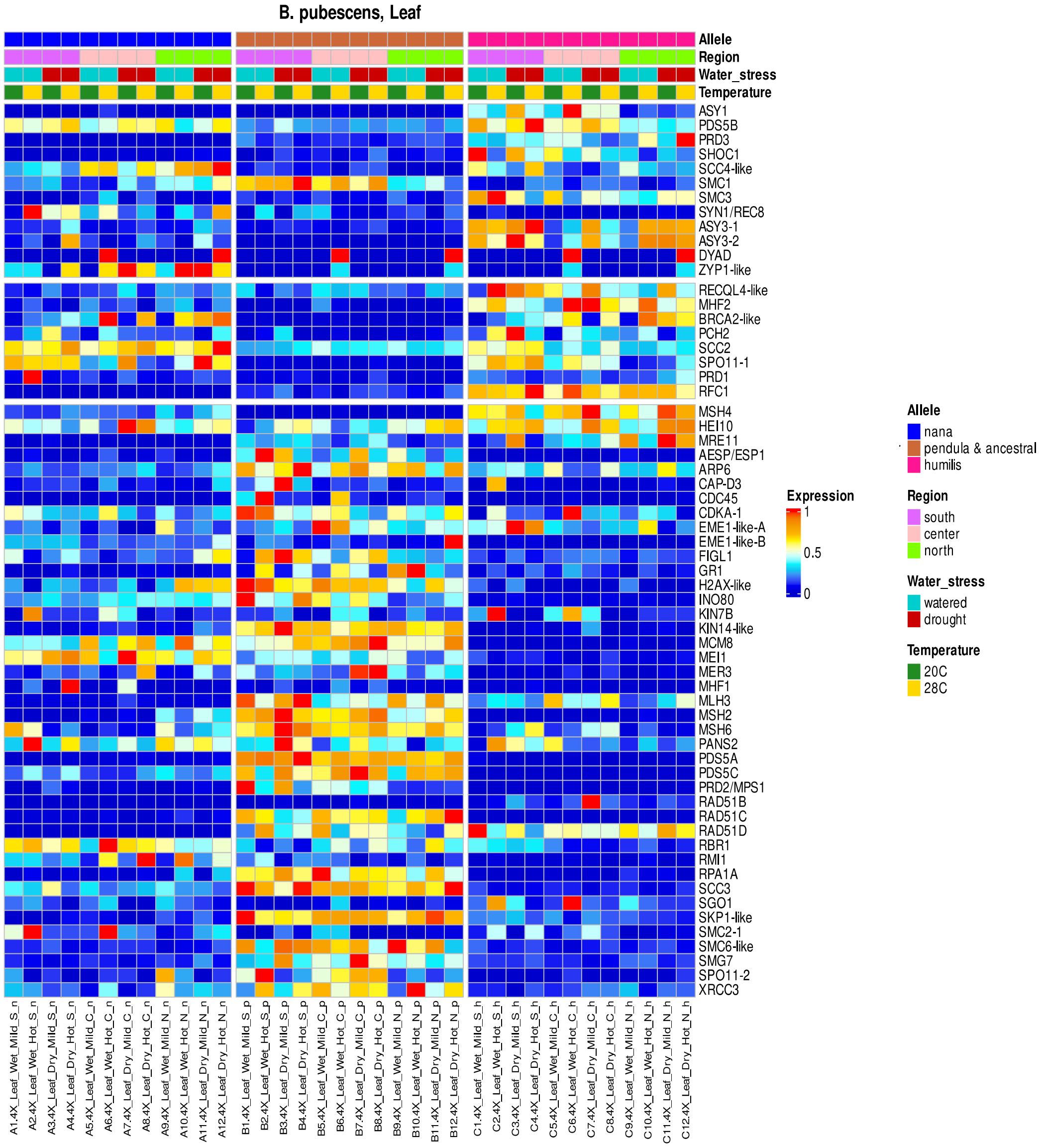
B. pubescens, Leaf
Allele
Region
Water_stress
Temperature
ASY1
PDS5B
PRD3
SHOC1
SCC4-like
SMC1
SMC3
SYN1/REC8
ASY3-1
ASY3-2
DYAD
ZYP1-like
RECQL4-like
MHF2
BRCA2-like
PCH2
SCC2
SPO11-1
PRD1
RFC1
MSH4
HEI10
MRE11
AESP/ESP1
ARP6
CAP-D3
CDC45
CDKA-1
EME1-like-A
EME1-like-B
FIGL1
GR1
H2AX-like
INO80
KIN7B
KIN14-like
MCM8
MEI1
MER3
MHF1
MLH3
MSH2
MSH6
PANS2
PDS5A
PDS5C
PRD2/MPS1
RAD51B
RAD51C
RAD51D
RBR1
RMI1
RPA1A
SCC3
SGO1
SKP1-like
SMC2-1
SMC6-like
SMG7
SPO11-2
XRCC3
A1.4X_Leaf_Wet_Mild_S_n
A2.4X_Leaf_Wet_Hot_S_n
A3.4X_Leaf_Dry_Mild_S_n
A4.4X_Leaf_Dry_Hot_S_n
A5.4X_Leaf_Wet_Mild_C_n
A6.4X_Leaf_Wet_Hot_C_n
A7.4X_Leaf_Dry_Mild_C_n
A8.4X_Leaf_Dry_Hot_C_n
A9.4X_Leaf_Wet_Mild_N_n
A10.4X_Leaf_Wet_Hot_N_n
A11.4X_Leaf_Dry_Mild_N_n
A12.4X_Leaf_Dry_Hot_N_n
B1.4X_Leaf_Wet_Mild_S_p
B2.4X_Leaf_Wet_Hot_S_p
B3.4X_Leaf_Dry_Mild_S_p
B4.4X_Leaf_Dry_Hot_S_p
B5.4X_Leaf_Wet_Mild_C_p
B6.4X_Leaf_Wet_Hot_C_p
B7.4X_Leaf_Dry_Mild_C_p
B8.4X_Leaf_Dry_Hot_C_p
B9.4X_Leaf_Wet_Mild_N_p
B10.4X_Leaf_Wet_Hot_N_p
B11.4X_Leaf_Dry_Mild_N_p
B12.4X_Leaf_Dry_Hot_N_p
C1.4X_Leaf_Wet_Mild_S_h
C2.4X_Leaf_Wet_Hot_S_h
C3.4X_Leaf_Dry_Mild_S_h
C4.4X_Leaf_Dry_Hot_S_h
C5.4X_Leaf_Wet_Mild_C_h
C6.4X_Leaf_Wet_Hot_C_h
C7.4X_Leaf_Dry_Mild_C_h
C8.4X_Leaf_Dry_Hot_C_h
C9.4X_Leaf_Wet_Mild_N_h
C10.4X_Leaf_Wet_Hot_N_h
C11.4X_Leaf_Dry_Mild_N_h
C12.4X_Leaf_Dry_Hot_N_h
Expression
1
0.5
0
Allele
nana
pendula & ancestral
humilis
Region
south
center
north
Water_stress
watered
drought
Temperature
20C
28C
